# Genomic evidence for global ocean plankton biogeography shaped by large-scale current systems

**DOI:** 10.1101/867739

**Authors:** Daniel J. Richter, Romain Watteaux, Thomas Vannier, Jade Leconte, Paul Frémont, Gabriel Reygondeau, Nicolas Maillet, Nicolas Henry, Gaëtan Benoit, Ophélie Da Silva, Tom O. Delmont, Antonio Fernàndez-Guerra, Samir Suweis, Romain Narci, Cédric Berney, Damien Eveillard, Frederick Gavory, Lionel Guidi, Karine Labadie, Eric Mahieu, Julie Poulain, Sarah Romac, Simon Roux, Céline Dimier, Stefanie Kandels, Marc Picheral, Sarah Searson, Tara Oceans Coordinators, Stéphane Pesant, Jean-Marc Aury, Jennifer R. Brum, Claire Lemaitre, Eric Pelletier, Peer Bork, Shinichi Sunagawa, Fabien Lombard, Lee Karp-Boss, Chris Bowler, Matthew B. Sullivan, Eric Karsenti, Mahendra Mariadassou, Ian Probert, Pierre Peterlongo, Patrick Wincker, Colomban de Vargas, Maurizio Ribera d’Alcalà, Daniele Iudicone, Olivier Jaillon

**Affiliations:** Institut de Biologia Evolutiva (CSIC-Universitat Pompeu Fabra), Passeig Marítim de la Barceloneta 37-49, 08003 Barcelona, Spain; Sorbonne Université, CNRS, Station Biologique de Roscoff, UMR 7144, ECOMAP, 29680 Roscoff, France; Stazione Zoologica Anton Dohrn, Villa Comunale, 80121 Naples, Italy; CEA, DAM, DIF, F - 91297 Arpajon Cedex, France; Aix Marseille Univ., Université de Toulon, CNRS, IRD, MIO UM 110,13288, Marseille, France; Génomique Métabolique, Genoscope, Institut de Biologie François Jacob, Commissariat à l’Énergie Atomique, CNRS, Université Evry, Université Paris-Saclay, Evry, France; Research Federation for the study of Global Ocean systems ecology and evolution, FR2O22/Tara GOsee, Paris, France; Changing Ocean Research Unit, Institute for the Oceans and Fisheries, University of British Columbia. Aquatic Ecosystems Research Lab. 2202 Main Mall. Vancouver, BC V6T 1Z4. Canada; Ecology and Evolutionary Biology, Yale University, New Haven, CT, USA; Institut Pasteur - Bioinformatics and Biostatistics Hub - C3BI, USR 3756 IP CNRS - Paris, France; INRIA/IRISA, Genscale team, UMR6074 IRISA CNRS/INRIA/Université de Rennes 1, Campus de Beaulieu, 35042, Rennes, France; Sorbonne Universités, CNRS, Laboratoire d’Oceanographie de Villefranche, LOV, F-06230 Villefranche-sur-Mer, France; Lundbeck Foundation GeoGenetics Centre, GLOBE Institute, University of Copenhagen, Øster Voldgade 5-7, 1350 Copenhagen K, Denmark; MARUM, Center for Marine Environmental Sciences, University of Bremen, Bremen, Germany; Max Planck Institute for Marine Microbiology, Celsiusstrasse 1, D-28359 Bremen, Germany; Dipartimento di Fisica e Astronomia ‘G. Galilei’ & CNISM, INFN, Università di Padova, Via Marzolo 8, 35131 Padova, Italy; MaIAGE, INRA, Université Paris-Saclay, 78350, Jouy-en-Josas, France; Université de Nantes, Centrale Nantes, CNRS, LS2N, F-44000 Nantes, France; Genoscope, Institut de biologie François-Jacob, Commissariat à l’Energie Atomique (CEA), Université Paris-Saclay, Evry, France; Sorbonne Universités, UPMC Université Paris 06, CNRS, Laboratoire d’oceanographie de Villefranche (LOV), Observatoire Océanologique, 06230 Villefranche-sur-Mer, France; Department of Oceanography, University of Hawaii, Honolulu, Hawaii 96822, USA; Department of Microbiology, The Ohio State University, Columbus, OH 43214, USA; Ecole Normale Supérieure, PSL Research University, Institut de Biologie de l’Ecole Normale Supérieure (IBENS), CNRS UMR 8197, INSERM U1024, 46 rue d’Ulm, F-75005 Paris, France; Structural and Computational Biology, European Molecular Biology Laboratory, Meyerhofstr. 1, 69117 Heidelberg, Germany; Directors’ Research European Molecular Biology Laboratory Meyerhofstr. 1 69117 Heidelberg Germany; Sorbonne Universités, UPMC Univ Paris 06, UMR 7093 LOV, F-75005, Paris, France; CNRS, UMR 7093 LOV, F-75005, Paris, France; PANGAEA, Data Publisher for Earth and Environmental Science, University of Bremen, Bremen, Germany; Department of Oceanography and Coastal Sciences, Louisiana State University, Baton Rouge, LA, 70808, USA; Max Delbrück Centre for Molecular Medicine, 13125 Berlin, Germany; Department of Bioinformatics, Biocenter, University of Würzburg, 97074 Würzburg, Germany; Institute of Microbiology, Department of Biology, ETH Zurich, Vladimir-Prelog-Weg 4, 8093 Zurich, Switzerland; Institut Universitaire de France (IUF), Paris, France; School of Marine Sciences, University of Maine, Orono, Maine 04469, USA; EMERGE Biology Integration Institute, The Ohio State University, Columbus, Ohio 43210, USA; Center of Microbiome Science, The Ohio State University, Columbus, Ohio 43210, USA; The Interdisciplinary Biophysics Graduate Program, The Ohio State University, Columbus, Ohio 43210, USA; Department of Civil, Environmental and Geodetic Engineering, The Ohio State University, Columbus OH 43214 USA; Department of Marine Biology and Oceanography, Institut de Ciències del Mar (ICM), CSIC, Barcelona, Spain; European Molecular Biology Laboratory, European Bioinformatics Institute (EMBL-EBI), Wellcome Trust Genome Campus, Hinxton, Cambridge CB10 1SD, United Kingdom; CNRS, UMR 7232, BIOM, Avenue Pierre Fabre, 66650 Banyuls-sur-Mer, France; Sorbonne Universités Paris 06, OOB UPMC, Avenue Pierre Fabre, 66650 Banyuls-sur-Mer, France; Aix Marseille Univ., Université de Toulon, CNRS, IRD, MIO UM 110, 13288, Marseille, France; Institute for Chemical Research, Kyoto University, Gokasho, Uji, Kyoto, 611-0011, Japan; Department of Microbiology and Immunology, Rega Institute, KU Leuven, Herestraat 49, 3000 Leuven, Belgium; VIB Center for Microbiology, Herestraat 49, 3000 Leuven, Belgium; CNRS, UMR 7009 Biodev, Observatoire Océanologique, F-06230 Villefranche-sur-mer, France; National Science Foundation, Arlington, VA 22230, USA; Bigelow Laboratory for Ocean Sciences East Boothbay, ME, USA; Laboratoire de Physique des Océans, UBO-IUEM, Place Copernic, 29820 Plouzané, France; Department of Geosciences, Laboratoire de Météorologie Dynamique (LMD), Ecole Normale Supérieure, 24 rue Lhomond, 75231 Paris Cedex 05, France

**Author notes:** equal contributions (Daniel J. Richter, Romain Watteaux, Thomas Vannier, Daniele Iudicone, Olivier Jaillon).

## Abstract

Biogeographical studies have traditionally focused on readily visible organisms, but recent technological advances are enabling analyses of the large-scale distribution of microscopic organisms, whose biogeographical patterns have long been debated. Here we assessed the global structure of plankton geography and its relation to the biological, chemical and physical context of the ocean (the ‘seascape’) by analyzing metagenomes of plankton communities sampled across oceans during the *Tara* Oceans expedition, in light of environmental data and ocean current transport. Using a consistent approach across organismal sizes that provides unprecedented resolution to measure changes in genomic composition between communities, we report a pan-ocean, size-dependent plankton biogeography overlying regional heterogeneity. We found robust evidence for a basin-scale impact of transport by ocean currents on plankton biogeography, and on a characteristic timescale of community dynamics going beyond simple seasonality or life history transitions of plankton.

## Main Text

Plankton communities are constantly on the move, transported by ocean currents^1^. Transport involves both advection and mixing. While being advected by currents, plankton can be influenced by multiple processes, both physico-chemical (fluxes of heat, light and nutrients^2^) and biological (species interactions, life cycles, behavior, acclimation/adaptation^3,4^), which act across various spatial and temporal scales. In turn, plankton impact seawater physico-chemistry while they are being advected^2^. The community composition and biogeochemical properties of a water mass at a given site are also partially dependent on its history of mixing with neighboring water masses during transport. These intertwined processes occurring along transport by currents form the pelagic seascape^5^ (Supplementary Fig. 1a). Due to logistical and analytical constraints, previous studies on plankton distribution have tended to be geographically or taxonomically restricted^6^–^10^, to focus on individual factors such as nutrient or light availability^11,12^, or have investigated the influence of transport on specific nutrients^13^ or types of planktonic organisms^14^–^16^. We set out to test for the first time at genomic resolution the hypotheses that a global-scale plankton biogeography exists and that it is closely linked to transport via large-scale ocean currents. To do this, we integrated metagenomic data from samples collected during the world-wide *Tara* Oceans expedition^17^ with *in situ* and satellite environmental metadata and large-scale ocean circulation simulations. The use of DNA as a primary proxy for global plankton diversity has several important advantages over classical morphology-based analyses, notably because methods can be standardized and applied across the entire range of plankton sizes, from viruses through prokaryotes and protists to animals.

DNA sequence data was obtained from samples collected at 113 world-wide stations during the *Tara* Oceans expedition, including from up to six organismal size fractions: one virus-enriched (0-0.22 μm)^8^, one prokaryote-enriched (either 0.22-1.6 or 0.22-3 μm)^18^, and four eukaryote-enriched (0.8-5 μm, 5-20 μm, 20-180 μm and 180-2000 μm^19^; Supplementary Fig. 1b). We analyzed 24.2 terabases of metagenomic sequence reads and 738 million eukaryotic 18S V9 ribosomal DNA marker sequences (Supplementary Table 1), complementing previously described *Tara* Oceans data^8,18,19^. We used metagenomic data and Operational Taxonomic Units (OTUs, representing groups of genetically related organisms) independently to compute pairwise comparisons of plankton community dissimilarity (as proxies for β-diversity). Metagenomic dissimilarity highlighted, at species and sub-species resolution, differences in the genomic identity of organisms between stations. Our metagenomic sampling resulted in pairwise metagenomic dissimilarities that likely represent an overestimate of β-diversity (Supplementary Information 1). However, we applied an identical procedure to compute metagenomic dissimilarity for all size fractions (correlations among fractions ranged from Spearman’s *ρ* 0.6 to 0.9, p ≤ 10^-4^, Supplementary Fig. 2). The more thoroughly sampled OTU dissimilarity, in contrast, incorporated more numerous rare taxa within the plankton, but at genus or higher-level taxonomic resolution^19^. Metagenomic and OTU dissimilarities were correlated for all size fractions (Spearman’s *ρ* 0.53 to 0.97, p ≤ 10^-4^, Supplementary Fig. 2), indicating that both proxies, although characterized by different sampling levels and taxonomic resolution, provided coherent and complementary estimates of β-diversity (Supplementary Information 1). We performed subsequent analyses using both measures, which produced consistent results. The taxonomic composition of these *Tara* Oceans samples, not discussed here, is instead presented in a parallel analysis^20^ of the spatial dynamics of planktonic eukaryotes, based on the same environmental data and large-scale ocean circulation simulations.

We focus on analyses of metagenomic dissimilarity here, with accompanying results for OTU dissimilarity presented in Supplementary Figures, and validation by comparison to abundance differences among metagenome-assembled genomes^21^ and to more traditional imaging data presented independently below.

Globally, we observed significant metagenomic dissimilarities between sampled stations (including adjacent sites) across all size fractions (Supplementary Fig. 3a, Supplementary Information 1). The resulting portrait is of a heterogeneous oceanic ecosystem at all scales separating *Tara* Oceans sampling sites (even those separated by only a few kilometers), dominated by a small number of abundant and cosmopolitan taxa, with a much larger number of less abundant taxa found at fewer sampling sites (Supplementary Fig. 3b-e), corroborating other studies^19,20^.

Overlying this heterogeneity, we found robust evidence for the existence of large-scale biogeographical patterns within all plankton size classes using two complementary analyses of dissimilarity among samples (Fig. 1a, Supplementary Fig. 4a-f, Supplementary Fig. 5, Supplementary Information 2). First, we grouped metagenomic samples within each size fraction into ‘genomic provinces’ via hierarchical clustering (Supplementary Fig. 6). Second, we derived colors for each sample based on a principal coordinates analysis (PCoA-RGB; see Methods) in order to visualize transitions in community composition within and between genomic provinces. Most genomic provinces were composed of large-scale geographically contiguous stations (consistent with previous studies documenting patterns in plankton biogeography^6-9^) with some independent distant samples (Fig. 1a, Supplementary Fig. 4a-f). Genomic provinces of smaller plankton (viruses, bacteria and eukaryotes <20 μm) tended to be limited to a single ocean basin and to approximately correspond to Longhurst biogeochemical provinces^11^ (Supplementary Fig. 4a-d; Supplementary Information 3). In contrast, provinces of larger plankton (micro- and meso-plankton, >20 μm) spanned multiple basins (Supplementary Fig. 4e-f, Supplementary Information 4).

**Figure 1.**
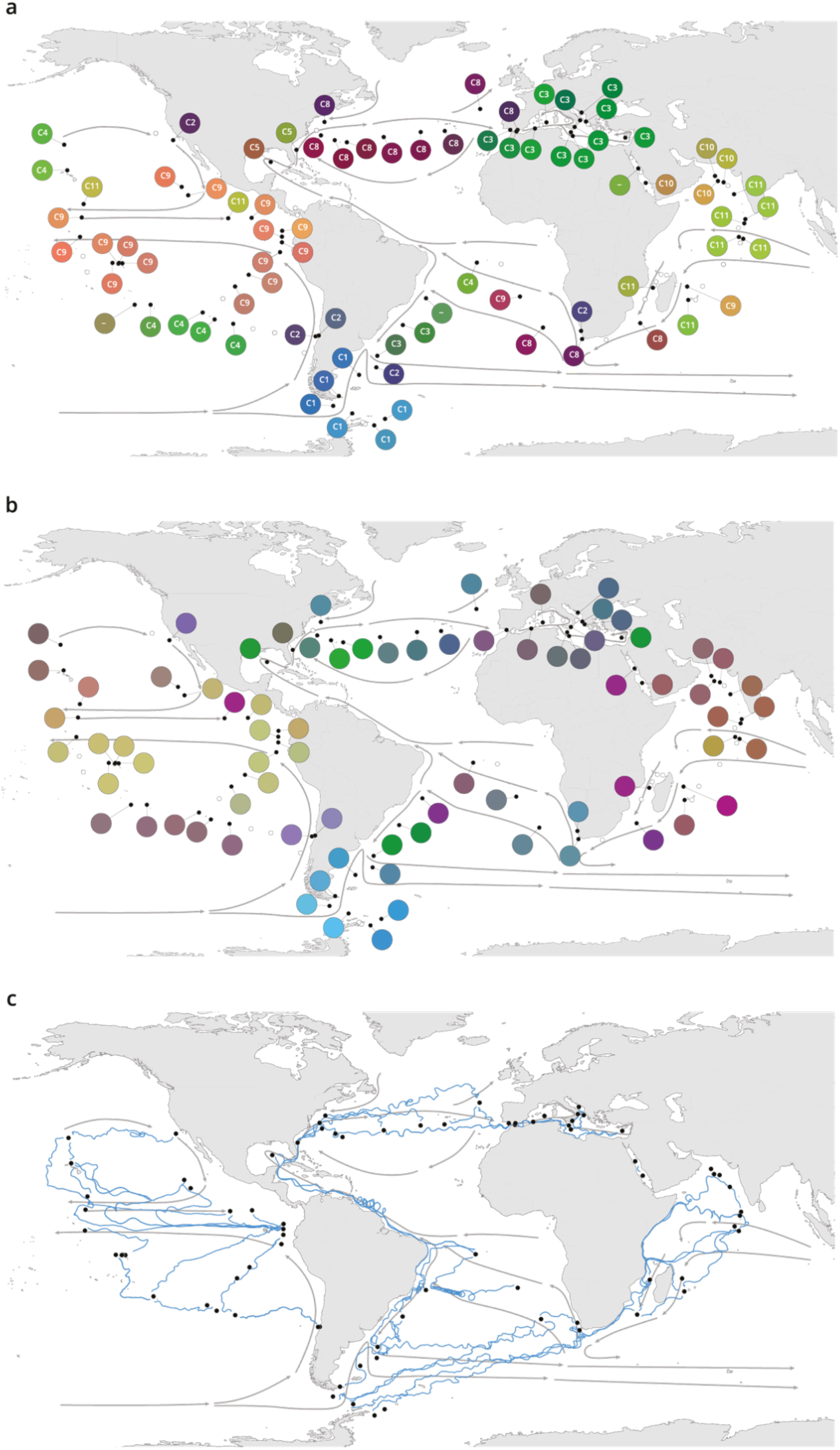
Plankton biogeography, environmental variation and ocean transport among *Tara* Oceans stations. Major currents are represented by solid arrows. **a**, Genomic provinces of *Tara* Oceans surface samples for the 0.8-5 μm size fraction, each labeled with a letter prefix (‘C’ represents the 0.8-5 μm size fraction) and a number; samples not assigned to a genomic province are labeled with ‘-’. Maps of all six size fractions and including DCM samples are available in Supplementary Fig. 4. Station colors are derived from an ordination of metagenomic dissimilarities; more dissimilar colors indicate more dissimilar communities (see Methods). **b**, Stations colored based on an ordination of temperature and the ratio of NO_3_ + NO_2_ to PO_4_ (replaced by 10^-6^ for 3 stations where the measurement of PO_4_ was 0) and of NO_3_ + NO_2_ to Fe. Colors do not correspond directly between maps; however, the geographical partitioning among stations is similar between the two maps. **c**, Simulated trajectories corresponding to the minimum travel time (T_min_) for pairs of stations (black dots) connected by T_min_ < 1.5 years. Directionality of trajectories is not represented.

These large-scale biogeographical patterns derived from metagenomes were linked to environmental parameters including nutrients and temperature. Seawater temperature was significantly different among genomic provinces for all plankton size classes (Kruskal-Wallis test, p < 10^-5^), corroborating previous results for prokaryotes^18^, whereas other environmental conditions were significantly different only with respect to specific size classes (Supplementary Fig. 7). The geography of combined nutrient and temperature variations resembled the biogeography of smaller plankton size classes (Fig. 1a-b, Supplementary Fig. 4a-d,h), whereas temperature alone more closely matched the distribution of larger plankton (Supplementary Fig. 4e,f,i), potentially reflecting different ecological constraints.

Many genomic provinces were spatially consistent with ocean basin-scale circulation patterns, such as western boundary currents or major subtropical gyres^22^ (Fig. 1a, Supplementary Fig. 4a-f), suggesting a particular role for large-scale surface transport (a core component of the seascape) in the emergence of spatial patterns of plankton community composition, as previously proposed^23^. We therefore investigated community metagenomic composition differences between sampled stations in light of the corresponding transit time, which has previously been suggested as the relevant factor for studying dispersal mechanisms^16^. We inferred the characteristic timescale of main transport paths between stations from trajectories computed with the physically well-constrained MITgcm ocean model (see Methods), which takes into account directionalities^1^ and meso-to large-scale circulation, potential dispersal barriers and mixing effects^24,25^. For this we used the minimum travel time^26^ (T_min_) between pairs of *Tara* stations. These trajectories corresponded to the dominant paths that transport the majority of water volume and its contents (e.g., heat, nutrients and plankton; Fig. 1c). For all plankton size classes, community composition differences between stations were significantly correlated to travel time (Supplementary Fig. 8).

Cumulative correlation values (correlations between metagenomic dissimilarity and T_min_ computed for an increasing range of T_min_) were maximal for pairs of stations separated by T_min_ <~1.5 years for all size classes, with correlation values (Spearman’s *ρ* 0.45 to 0.71 depending on size class, p ≤ 10^-4^; Fig. 2a, Supplementary Fig. 9) far exceeding those based on previous studies of morphological and/or metabarcode data^15^ or considering geographic distance rather than travel time^27^. These high correlations between metagenomic dissimilarity and T_min_ for travel times up to 1.5 years hence reveal measurable plankton community dynamics on time scales far longer than typical plankton growth rates or life cycles. In contrast, no such unimodal pattern was found for correlations between metagenomic dissimilarity and geographic distance (without traversing land; Supplementary Fig. 9f). Over the timescale <~1.5 years, which corresponds well with the average time to travel across a basin or gyre, the timescale of large-scale transport is therefore an appropriate framework for studying differences in plankton genomic community composition (Fig. 2b). The fact that simulated transport times and metagenomic dissimilarity were correlated despite a 3 year pan-season sampling campaign, which could be considered to weaken our inference, suggests instead that a large-scale impact of the seascape promotes the existence of a biogeographical structure at a large spatial scale that is resilient to seasonal or other smaller spatio-temporal variations (across all size fractions, genomic provinces consist of stations sampled over an average of 4.7 ± 2.8 different months and 2.7 ± 1.2 different seasons, adjusted for hemisphere). Consistent with our results, seasonal variations have previously been shown to have minor effects on the boundary positions of biogeochemical provinces based on satellite data, but not enough to affect the overall pattern of ocean regionalization^28^.

**Figure 2.**
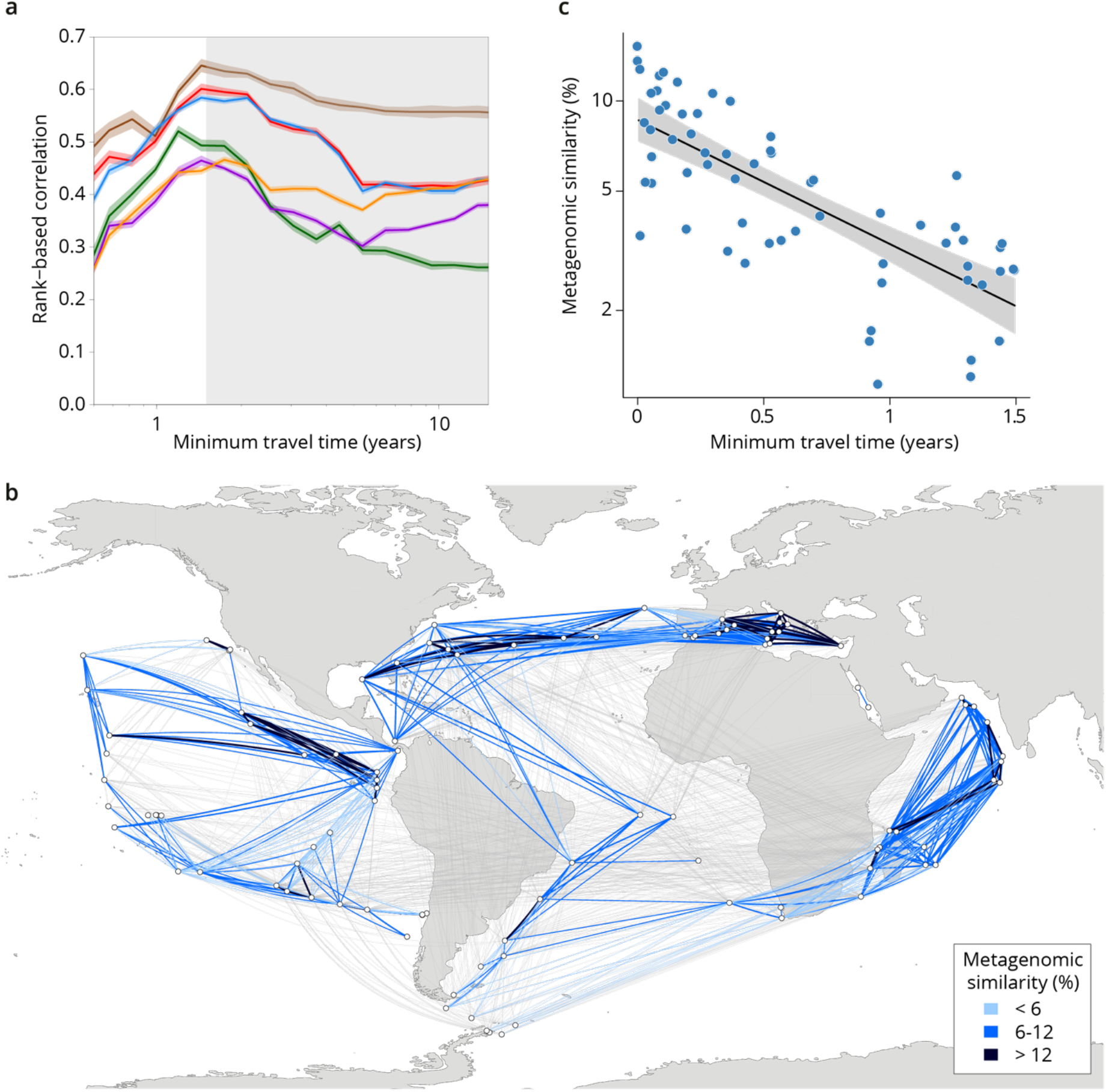
Metagenomic dissimilarity and travel time of plankton are maximally correlated up to ~1.5 years. **a**, Spearman rank-based correlation by size fraction between metagenomic dissimilarity and minimum travel time along ocean currents (T_min_) for pairs of *Tara* Oceans samples separated by a minimum travel time less than the value of T_min_ on the x axis. Brown line: 0-0.2 μm size fraction, red: 0.22-1.6/3 μm, blue: 0.8-5 μm, green: 5-20 μm, purple: 20-180 μm, orange: 180-2000 μm. Shaded colored areas represent 95% confidence intervals. T_min_ >1.5 years is shaded in grey. See plots for OTU dissimilarity in Supplementary Fig. 9. **b**, Pairs of *Tara* stations connected by T_min_ <1.5 years in blue/black and >1.5 years in grey. Shading reflects metagenomic similarity from the 0.8-5 μm size fraction. **c**, The relationship of metagenomic similarity to T_min_ with an exponential fit (black line, grey 95% CI), for pairs of surface samples in the 0.8-5 μm size fraction within the North Atlantic and Mediterranean current system (see map and plots for other size fractions and OTUs in Supplementary Fig. 10, and Supplementary Information 1 for a discussion of metagenomic similarity).

Differences in environmental conditions for pairs of stations also covaried (although less strongly) with transit time for T_min_ <~1.5 years (Fig. 3). This indicates that changes in environmental conditions and plankton community composition are concurrent along large-scale oceanic current systems. In our data, beyond ~1.5 years of transport, correlations of T_min_ with metagenomic dissimilarity decreased (Fig. 2a, Fig. 3, Supplementary Fig. 9a-e), meaning the signature of transport in generating large-scale diversity changes weakened and travel time therefore becomes a less appropriate context to study β-diversity. A similar trend was observed for the correlation between T_min_ and nutrient concentrations, whereas temperature, the gradients of which are mostly dictated by Earth-scale processes, remained well correlated for longer transit times (Fig. 3).

**Figure 3.**
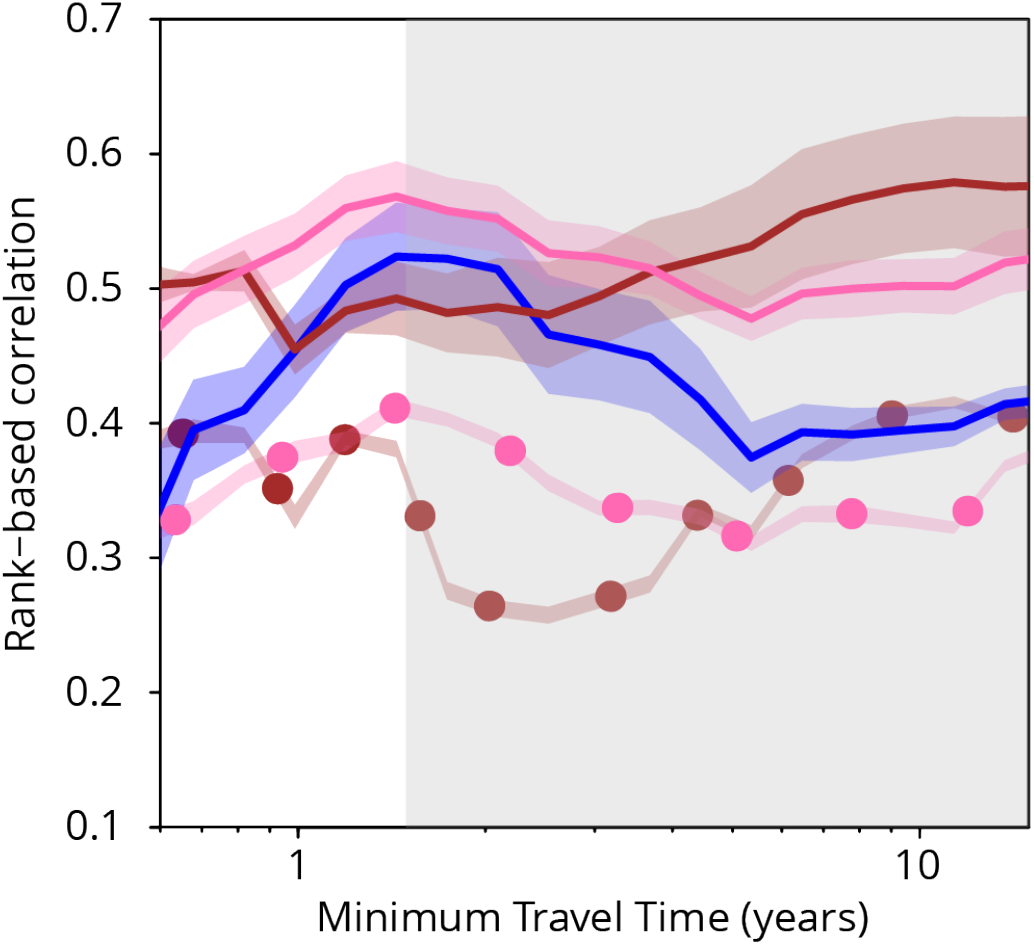
Plankton travel time, metagenomic dissimilarity and environmental differences show different temporal patterns of pairwise correlation. Spearman rank-based correlations between metagenomic dissimilarity and minimum travel time (T_min_, blue), metagenomic dissimilarity and differences in NO_3_ + NO_2_, PO_4_ and Fe (pink), metagenomic dissimilarity and differences in temperature (red), T_min_ and differences in NO_3_ + NO_2_, PO_4_ and Fe (pink, dashed), and T_min_ and differences in temperature (red, dashed) for pairs of *Tara* Oceans samples separated by a minimum travel time less than the value of T_min_ on the x axis. Shaded regions represent standard error of the mean. Correlations represent averages across four of six size fractions represented in Fig. 2a; the 0-0.2 μm and 5-20 μm size fractions are excluded due to a lack of samples at the global level. Individual size fractions, partial correlations, and correlations with OTU data are in Supplementary Fig. 9.

Together, these analyses suggest the existence in the seascape of biogeochemical continua stretched by currents on the basin scale with predictable, interlinked changes in environmental conditions and plankton community composition (Supplementary Information 5). It has previously been posited that transport could generate continuous transitions between niches based on physical processes^29^, but it was not anticipated that other aspects of the seascape would be implicated and that this would occur on the scale of ocean basins or larger. Moreover, beyond ~1.5 years, the correlation of metagenomic dissimilarity with differences in temperature increased while that with differences in nutrients decreased (Fig. 3, Supplementary Fig. 9a-e), although both of these correlations with metagenomic dissimilarity remained strong on these time scales. This might be related to distant *Tara* Oceans stations experiencing similar oceanographic phenomena (notably temperature), for example upwelling zones, producing generally similar environmental conditions.

The existence of a size-class dependent (smaller or larger than 20 μm) structure of plankton geography indicates that the continua that we observe vary among size fractions because of different reactions of organisms within the seascape, in agreement with a parallel survey based on taxonomic groups^20^. In the case of the North Atlantic current system (including the Mediterranean Sea), a simple exponential fit of metagenomic dissimilarity along T_min_ for T_min_ <~1.5 years (Fig. 2c) revealed that the smaller size classes (<20 μm) had a shorter metagenomic turnover time (ca. 1y) than larger plankton (ca. 2y) (Supplementary Fig. 10, Supplementary Information 6). At global geographical scales, the genomic provinces of small size classes, which are enriched in phytoplankton^18-20,30^, corresponded in our data with differences in environmental parameters such as nutrient levels (Fig. 1b, Supplementary Fig. 7) that are often constrained by regional oceanographic processes^31^. On the other hand, genomic provinces of larger plankton, enriched in heterotrophic and symbiotic organisms^19,20,30^, were less coupled with geochemical parameters and were more related to global scale gradients and circulation patterns, notably major latitudinal temperature zones or the separation between Atlantic and Indo-Pacific large-scale surface circulations (Supplementary Fig. 4e,f,i). These divergent effects were also evident in comparisons of metagenomic dissimilarity with variations in environmental conditions (Supplementary Fig. 9b). For smaller plankton, correlations with differences in nutrient concentrations were stronger for T_min_ up to ~1.5 years, but for larger plankton, correlations were stronger with temperature variations for T_min_ beyond ~1.5 years. Larger plankton are dominated by eukaryotes, often multicellular, with much longer life cycles, potentially leading to slower community turnover. Organisms with long life cycles, on the order of several months or years, can be transported through basins spanning multiple biogeochemical niches in which they may encounter strong environmental variability; this trend was also detected in a taxonomy-based analysis accounting for differences in both body size and ecology among groups^20^. As observed here, their biogeography is less affected by nutrient limitation and rather depends on large-scale temperature gradients among basins. This dependence may be linked to the known correlation between body size and organismal metabolic rate^32^. Conversely, variants within populations of organisms with short life cycles have the capacity to increase their relative abundance within restricted ecological niches to which they are adapted. This difference, detectable at genomic resolution, may not be picked up in analyses performed using biological traits with less resolution. These results indicate a significant size-based decoupling within planktonic food webs. For example, large size predators will encounter different prey when transiting through the genomic provinces of small sized organisms (see Supplementary Information 4).

We compared our analyses of metagenomic data to those based on more traditional zooplankton imaging data collected for the same *Tara* Oceans samples. β-diversity calculated from zooplankton imaging was correlated with metagenomic dissimilarity (Spearman’s ρ between 0.32 and 0.60; Supplementary Fig. 2), indicating that the two data sources provide concordant measurements of variation in plankton community composition. However, correlations with ocean transport time were far weaker for zooplankton imaging data than for metagenomic data from all organismal size fractions (Supplementary Fig. 9), to the extent that we were not able to calculate community turnover times based on imaging data from the same set of stations using an exponential fit. We interpret this as being a result of the expected significantly lower resolution in imaging data as compared to metagenomic data (a similar difference of resolution in OTU data versus metagenomic data is discussed in Supplementary Information 1). Finally, we also confirmed our metagenome sequence read comparison-based results by comparing them to β-diversity among sampling sites using a collection of metagenome-assembled genomes (MAGs), which are likely to represent the most abundant genomes, from the 20-180 μm size fraction (the size fraction in which the largest proportion of metagenomic reads were mapped to MAGs, 18.4%)^21^. Metagenomic and MAG β-diversity were highly correlated (Spearman’s ρ 0.94) and consequently they displayed similar biogeographical patterns (Supplementary Fig. 4e,g).

In this study, we provide genomic evidence for an organism-size-dependent global-scale plankton biogeography shaped by ocean currents. Using analysis of standardized metagenomic data, we reveal that the integration of seascape physical, chemical and biological processes over time and space produces a quasi-stationary biological partitioning of the oceans that supersedes short-term variability and seasonal cycles, ultimately generating global biogeographical patterns. Future studies both on smaller spatio-temporal scales and on the global-scale constraints and influences on the seascape itself (i.e., the three-dimensional topology of the oceans) could lead to a more detailed understanding of plankton dynamics. Overall, our work shows that studies of the dynamics of plankton communities must consider the critical influence of ocean currents in stretching and altering, on the scale of basins, the distribution of both planktonic organisms and the physico-chemical nature of the water mass in which they reside. We also demonstrate that the combination of ocean circulation modeling with the use of metagenomic DNA as a tracer of plankton communities provides a resolution above the minimum necessary for assessing the role of transport in community turnover over time and space. The planktonic ecosystem is fundamentally different in many ways from other major planetary ecosystems and this study provides a basis to understand and potentially predict the structuring of the ocean ecosystem in a scenario of rapid environmental and current system changes^30,33,34^.

## Methods

### Sampling, sequencing and environmental parameters

Sampling, size fractionation, measurement of environmental parameters and associated metadata, DNA extraction and metagenomic sequencing were conducted as described previously^35,36^. Samples were collected at 113 *Tara* Oceans stations for up to six size fractions (0-0.2, 0.22-1.6/3, 0.8-5, 5-20, 20-180, 180-2000 μm; Supplementary Fig. 1b; Supplementary Table 1) and two depths (subsurface and deep chlorophyll maximum (DCM)). The prokaryote-enriched size fraction was collected either a 0.22-1.6 μm or 0.22-3 μm filter^18,35^. For technical reasons, not all size fractions were sequenced for all stations (see Supplementary Information 7 for a summary of why this does not affect our principal conclusions).

We used physico-chemical data measured *in situ* during the *Tara* Oceans expedition (depth of sampling, temperature, chlorophyll *a*, phosphate, nitrate + nitrite concentrations), supplemented with simulated values for iron and ammonium (using the MITgcm Darwin model described below in “Ocean circulation simulations”), day length, and 8-day averages calculated for photosynthetically active radiation (PAR) in surface waters (AMODIS, https://modis.gsfc.nasa.gov). In order to obtain PAR values at the deep chlorophyll maximum, we used the following formula^37^:

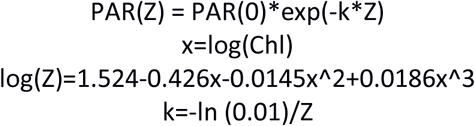

in which k is the attenuation coefficient, and Z is the depth of the DCM (in meters). Other data, such as silicate and the (nitrate + nitrite)/phosphate ratio, were extracted from the World Ocean Atlas 2013 (WOA13 version 2, https://www.nodc.noaa.gov/OC5/woa13/), by retrieving the annual mean values at the closest available geographical coordinates and depths to *Tara* sampling stations. For temperature and nitrate + nitrite, we calculated seasonality indexes (SI) from monthly WOA13 data. For each sample, the index is the annual variation of the parameter (max - min) at this location divided by the highest variation value among all samples.

A list of samples, metagenomic and metabarcode sequencing information and associated environmental data is available in Supplementary Tables 1-2.

### Calculation of metagenomic community dissimilarity

Metagenomic community distance between pairs of samples was estimated using whole shotgun metagenomes for all six size fractions. We used a metagenomic comparison method (Simka^38^) that computes standard ecological distances by replacing species counts by counts of DNA sequence k-mers (segments of length *k*). K-mers of 31 base pairs (bp) derived from the first 100 million reads sequenced in each sample (or the first 30 million reads for the 0-0.2 μm size fraction) were used to compute a similarity measure between all pairs of samples within each organismal size fraction. Based on a benchmark of Simka, we selected 100 million reads per sample (or 30 million for the 0-0.2 μm fraction) because increasing this number did not produce a qualitatively different set of results, and to ensure that the same number of reads were used in each pairwise comparison within a size fraction. Nearly all samples in our data set had at least 100 million reads (or at least 30 million for the 0-0.2 μm fraction; Supplementary Table 1).

We estimated β-diversity for metagenomic reads with the following equation within Simka:

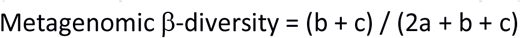

Where a is the number of distinct k-mers shared between two samples, and b and c are the number of distinct k-mers specific to each sample. We represented the distance between each pair of samples on a heatmap using the heatmap.2 function of the R-package^39^ gplots_2.17.0^40^. The dissimilarity matrices we produced for each plankton size fraction (on a scale of 0 = identical to 100 = completely dissimilar) are available as Supplementary Tables 3-8.

### Calculation of OTU-based community dissimilarity

Within the 0-0.2 μm size fraction, we used previously published viral populations (equivalent to OTUs)^41^ and viral clusters (analogous to higher taxonomic levels)^8^ based on clustering of protein content. For the 0.22-1.6/3 μm size fraction, we used previously derived miTAGs based on metagenomic matches to 16S ribosomal DNA loci and processed them as described^18^. For the four eukaryotic size fractions, we added additional samples to a previously published *Tara* Oceans metabarcoding data set and processed them using the same methods^19^ (also described at DOI: 10.5281/zenodo.15600).

We calculated OTU-based community dissimilarity for all size fractions as the Jaccard index based on presence/absence data using the vegdist function implemented in vegan 2.4-0^42^ in the software package R. The dissimilarity matrices we produced for each plankton size fraction (on a scale of 0 = identical to 100 = completely dissimilar) are available as Supplementary Tables 9-14.

### Calculating distances of environmental parameters

We calculated Euclidean distances^43^ for physico-chemical parameters. Each were scaled individually to have a mean of 0 and a variance of 1 and thus to contribute equally to the distances. Then the Euclidean distance between two stations i and j for parameters P was computed as follows:

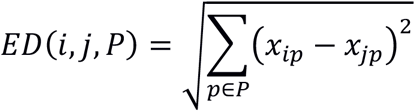

### RGB encoding of environmental positions

We color-coded the position of stations in environmental space for Fig. 1b and Supplementary Fig. 4h as follows. First, environmental variables were power-transformed using the Box-Cox transformation to have Gaussian-like distributions to mitigate the effect of outliers and scaled to have zero mean and unit variance. We then performed a principal component analysis (PCA) with the R command prcomp from the package stats 3.2.1^39^ on the matrix of transformed environmental variables and kept only the first 3 principal components. Finally, we rescaled the scores in each component to have unit variance and decorrelated them using the Mahalanobis transformation. Each component was mapped to a color channel (red, green or blue) and the channels were combined to attribute a single composite color to each station. The components (x, y, z) were mapped to color channel values (r, g, b) between 0 and 255 as r = 128 * (1 + x / max(abs(x)), and similarly for g and b. This map ensures that the global dispersion is equally distributed across the three components and composite colors span the whole color space.

### Definition of genomic provinces

We used a hierarchical clustering method on the metagenomic pairwise dissimilarities produced by Simka for all surface and DCM samples, and multiscale bootstrap resampling for assessing the uncertainty in hierarchical cluster analysis. We focused on metagenomic dissimilarity due to its higher resolution, and confirmed that the patterns found in metagenomic data were consistent when using OTU data (Supplementary Fig. 5). We used UPGMA (Unweighted Pair-Group Method using Arithmetic averages) clustering, as it has been shown to have the best performance to describe clustering of regions for organismal biogeography^44^. The R-package pvclust_1.3-2^45^, with average linkage clustering and 1,000 bootstrap replications, was used to construct dendrograms with the approximately unbiased p-value for each cluster (Supplementary Fig. 6). Because the number of genomic provinces by size fraction was not known *a priori*, we applied a combination of visualization and statistical methods to compare and determine the consistency within clusters of samples. First, the silhouette method^46^ was used to measure how similar a sample was within its own cluster compared to other clusters using the R package cluster_2.0.1^47^. The Silhouette Coefficient *s* for a single sample is given as:

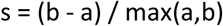

Where *a* is the mean distance between *a* sample and all other points in the same class and *b* is the mean distance between *a* sample and all other points in the next nearest cluster. We used the value of *s*, in addition to bootstrap values, to partition each tree into genomic provinces (see Supplementary Information 2 for further details on statistical validation of genomic provinces). Additionally, we used the Radial Reingold-Tilford Tree representation from the JavaScript library D3.js (https://d3js.org/)^48^ to visualize sample partitions from the dendrogram. Single samples were not considered as genomic provinces.

In a complementary approach, we performed a principal coordinates analysis (PCoA) with the R command cmdscale (eig = TRUE, add = TRUE) from the package stats 3.2.1^39^ on the matrices of pairwise metagenomic dissimilarities calculated by Simka (or OTU dissimilarity measured with the Jaccard index) within each size fraction and kept only the first 3 principal coordinates. We then converted those coordinates to a color using the RGB encoding described above, with one modification: scaling factors λ_r_, λ_g_ and λ_b_ were calculated as the ratios of the second and third eigenvalues to the first (dominant) eigenvalue to ensure that the dispersion of stations along each color channel reproduced the dispersion of the stations along the corresponding principal component (the ratio for the color corresponding to the dominant eigenvalue is 1). The components (x, y, z) were then mapped to color channel values (r, g, b) between 0 and 255 as r = 128 * (1 + λ_c_x / max(abs(x)), where λ_c_ is the ratio of the eigenvalue of color c to the dominant eigenvalue.

We represented number and PCoA-RGB color of genomic provinces for each sample on a world map (Fig. 1, Supplementary Fig. 4a-f) generated with the R packages maps_3.0.0.2^49^, mapproj 1.2-4^50^, gplots_2.17.0^40^ and mapplots_1.5^51^. We also plotted phosphate and temperature (Supplementary Fig. 4a-f) obtained from the *Csiro Atlas of Regional Seas* (CARS2009, http://www.cmar.csiro.au/cars) using the phosphate_cars2009.nc and temprerature_cars2009a.nc files and the R package RNetCDF^52^.

### Comparison of genomic provinces to previous ocean divisions

To evaluate the spatial similarity between the clusters obtained in our study for each size fraction and previous biogeographic divisions, we performed an analysis of similarity (ANOSIM, Fathom toolbox, matlab^®^). First, we collected coordinates for three spatial divisions at a resolution of 0.5° x 0.5°: biomes, biogeochemical provinces (BGCPs)^11,28^ and objective global ocean biogeographic provinces (OGOBPs)^53^. Second, we assigned *Tara* Oceans stations to biomes, BGCPs, and OGOBPs based on their GPS coordinates. Third, for each size fraction we performed an ANOSIM with the metagenomic dissimilarity matrix calculated by Simka, using biogeographic clusters (biome, BGCP, OGOBP) as group membership for each station. Each ANOSIM was bootstrapped 1,000 times to evaluate the interval of confidence around the strength of the relationships we detected (Supplementary Fig. 4a-f).

### Environmental differences among genomic provinces

For each size fraction, we tested which environmental parameters significantly discriminated among genomic provinces (Supplementary Fig. 7). A total of 12 parameters characterizing each sample, grouped by genomic provinces, were evaluated with a Kruskal-Wallis test within each size fraction with a significance threshold of p < 10^-5^. Selected parameters for each size fraction were then used to perform a principal components analysis of the samples using the R package vegan_1.17-11^42^. Samples were plotted with the same PCoA-RGB colors used in the genomic province maps above and each genomic province surrounded by a grey polygon. In analyses where Southern Ocean (including Antarctic) stations were considered independently from other stations, the following were considered Southern Ocean stations: 82, 83, 84, 85, 86, 87, 88, 89.

### Ocean circulation simulations

We derived travel times from the MITgcm Darwin simulation^54^ based on an optimized global ocean circulation model from the ECCO2 group^55^. The horizontal resolution of the model was approximately 18 km, with 1,103,735 total ocean cells. We ran the model for six continuous years in order to smooth anomalies that might occur during any single year. We used surface velocity simulation data to compute trajectories of floats originating in ocean cells containing all *Tara* Oceans stations, and applied the following stitching procedure to generate a large number of trajectories for each initial position. (The use of surface velocity data implies that Ekman transport also influences trajectories within the simulation.)

First, we precomputed a set of monthly trajectories: for each of the 72 months in the dataset, we released floats in every ocean cell of the model grid and simulated transport for one month. We used a fourth-order Runge-Kutta method with trilinearly interpolated velocities and a diffusion of 100 m^2^/s. Second, following previous studies^14^, we stitched together monthly trajectories to create 10,000 year trajectories: for each float released within a 200 km radius of a *Tara* station, we constructed 1,000 trajectories, each 10,000 years long. To avoid seasonal effects, we began by selecting a random starting month. We followed the trajectory of a float released within that month to the grid cell containing its end point at the end of the month. Next, we randomly selected a trajectory starting on the following month (e.g., February would follow January) from that grid cell, and repeated until reaching a 10,000 year trajectory.

We searched the resulting 50.8 million trajectories for those that connected pairs of *Tara* Oceans stations. To ensure robustness of our results, we only included pairs of stations that were connected by more than 1,000 trajectories. For each pair of stations, T_min_ was defined as the minimum travel time of all trajectories (if any) connecting the two stations. The travel time matrix we produced (measured in years) is available as Supplementary Table 15. Standard minimum geographic distance without traversing land^56^ is available as Supplementary Table 16.

### Correlations of β-diversity, T_min_ and environmental parameters

We excluded stations that were not from open ocean locations from correlation analyses to avoid sites impacted by coastal processes (those numbered 54, 61, 62, 79, 113, 114, 115, 116, 117, 118, 119, 120, and 121). In analyses where Southern Ocean (including Antarctic) stations were considered independently from other stations, the following were considered Southern Ocean (including Antarctic) stations: 82, 83, 84, 85, 86, 87, 88, 89. We calculated rank-based Spearman correlations between β-diversity, T_min_ and environmental parameters (either differences in temperature or the Euclidean distance composed of differences in NO_3_ + NO_2_, PO_4_ and Fe, see above) for surface samples with a Mantel test with 1,000 permutations and a nominal significance threshold of p < 0.01. For the correlations presented in Fig. 2a, Fig. 3 and Supplementary Fig. 9 correlation values were derived from pairs of stations connected by T_min_ up to the value on the x-axis. We calculated partial correlations of metagenomic and OTU dissimilarity and T_min_ by controlling for differences in temperature and for differences in nutrient concentrations, and partial correlations of dissimilarity with temperature or nutrient variation by controlling for T_min_.

### Community turnover in the North Atlantic

*Tara* Oceans stations numbered 72, 76, 142, 143, 144, and all stations from 146 to 151 were located along the main current system connecting South Atlantic and North Atlantic oceans and continuing to the strait of Gibraltar. In addition, we included stations 4, 7, 18, and 30 located on the main current system in the Mediterranean Sea (Supplementary Fig. 10). As the *Tara* Oceans samples within the subtropical gyre of the North Atlantic and in the Mediterranean Sea were all collected in winter, seasonal variations should not play a role in the variability in community composition that we observed (see Supplementary Table 2). We calculated genomic e-folding times (the time after which the detected genomic similarity between plankton communities changes by 63%) over scales from months to years based on an exponential fit of metagenomic dissimilarity to T_min_ with the form y = C_0_ e^-x/τ^ (where C_0_ is a constant and τ the folding time). Exponential fits for size fractions 0-0.2 μm and 5-20 μm were not calculated due to an insufficient number of sampled stations in the North Atlantic (Supplementary Information 6).

The synthetic map (Supplementary Fig. 10a) was generated with the R packages maps_3.0.0.2, mapproj 1.2.4, gplots_2.17.0 and mapplots_1.5. We derived dynamic sea surface height from the *Csiro Atlas of Regional Seas* (CARS2009, http://www.cmar.csiro.au/cars) using the hgt2000_cars2009a.nc file and plotted with the R package RNetCDF.

### Imaging methods

Plankton were also collected using WP2 (200 μm mesh) nets, using vertical tows (0-100 m) and preserved with borax-buffered formaldehyde. Taxonomic classification was performed using the ZooScan imaging system^57^ and identified with an automatic recognition algorithm to the finest possible taxonomic resolution using Ecotaxa^58^. The resulting identifications were manually visualized by taxonomic specialists and either validated or corrected. Resolution of the taxonomic identifications depended on morphological heterogeneity within taxonomic groups. Hence, identifications reached different taxonomic levels, from species to phylum, and most of them reached family level. All images and their taxonomic assignation are accessible within Ecotaxa (https://ecotaxa.obs-vlfr.fr/prj/377). Since all genomic data were collected during day-time, we restricted our analysis on day-collected samples. We also discarded non-living objects in our analyses. We estimated β-diversity by calculating Bray-Curtis dissimilarities between pairs of stations based on the relative abundances of each annotated taxonomic unit. Bray-Curtis dissimilarities are available as Supplementary Table 17.

### Metagenome-assembled genomes (MAGs) analysis

MAG relative abundances in metagenomic samples were retrieved from Delmont *et al*^21^. β-diversity was estimated by calculating the Bray-Curtis dissimilarities between pairs of stations based on the relative abundances of each of the 713 MAGs calculated by read mapping in the metagenomes of size fraction 20-180 μm (the size fraction in which MAGs recruit the largest relative share of all reads). We represented PCoA-RGB color of the Bray-Curtis dissimilarity matrix for each sample on a world map (Supplementary Fig. 4g) following the methodology described above. The Spearman ρ correlation coefficient was calculated between MAG-based β-diversity and metagenomic based β-diversity from the size fraction 20-180 μm. MAG-derived Bray-Curtis dissimilarities for the 20-180 μm size fraction are available as Supplementary Table 18.

## Acknowledgements

We acknowledge Oliver Jahn and M. J. Follows for providing numerical simulations of particle trajectories from *Tara* Oceans stations. We thank the commitment of the following people and sponsors who made this expedition possible: CNRS (in particular Groupement de Recherche GDR3280), European Molecular Biology Laboratory (EMBL), Genoscope/CEA, the French Government ‘Investissement d’Avenir’ programs OCEANOMICS (ANR-11-BTBR-0008) and FRANCE GENOMIQUE (ANR-10-INBS-09), Fund for Scientific Research - Flanders, VIB, Stazione Zoologica Anton Dohrn, UNIMIB, MEMO LIFE (ANR-10-LABX-54), Paris Sciences et Lettres (PSL) Research University (ANR-11-IDEX-0001-02), ANR (projects POSEIDON/ANR-09-BLAN-0348, PROMETHEUS/ANR-09-PCS-GENM-217, MAPPI/ANR-2010-COSI-004, TARA-GIRUS/ANR-09-PCS-GENM-218, HYDROGEN/ANR-14-CE23-0001), EU FP7 MicroB3/No. 287589, US NSF grant DEB-1031049, FWO, BIO5, Biosphere 2, Agnès b., the Veolia Environment Foundation, Région Bretagne, World Courier, Illumina, Cap L’Orient, the EDF Foundation EDF Diversiterre, FRB, the Prince Albert II de Monaco Foundation, Etienne Bourgois, the *Tara* schooner and its captain and crew. We thank MERCATOR-CORIOLIS and ACRI-ST for providing daily satellite data during the expedition. The bulk of genomic computations were performed using the Airain HPC machine provided through GENCI-[TGCC/CINES/IDRIS] (grants t2011076389, t2012076389, t2013036389, t2014036389, t2015036389 and t2016036389). We are also grateful to the French Ministry of Foreign Affairs for supporting the expedition and to the countries who granted us sampling permissions. *Tara* Oceans would not exist without continuous support from 23 institutes (http://oceans.taraexpeditions.org).

DJR was supported by postdoctoral fellowships from the Conseil Régional de Bretagne and from the Beatriu de Pinós programme of the Government of Catalonia’s Secretariat for Universities and Research of the Ministry of Economy and Knowledge. RW, DI and MRd’A were supported by the Italian Flagship Project RITMARE and Premiale MIUR NEMO. MBS was supported by US NSF grants OCE-1536989 and OCE-1829831, grant #3709 from the Gordon and Betty Moore Foundation, and HPC support from the Ohio Super Computer.

We also acknowledge Stéphane Audic for assistance with metabarcoding analyses, C. Scarpelli for support in high-performance computing, Mathieu Raffinot and Dominique Lavenier for discussions on sequence comparison algorithms, Samuel Chaffron for help with sample contextual data, Noan Le Bescot (Ternog Design) for assistance in preparing figures, and Marion Gehlen. We thank all members of the *Tara* Oceans consortium for maintaining a creative environment and for their constructive criticism.

## Author Contributions

DI, OJ, CdV, and PW designed and directed the study. IP, DJR, RW, OJ, DI, MRd’A, TV and CdV wrote the manuscript. TV, GB, NM, PP, CL and OJ designed and computed pairwise metagenomic comparisons. TV, DJR, RW, JL and PF performed the analyses of genomic data with substantial input from MRd’A, DI, OJ and PW. RW, DI, TV, PF and DJR analyzed ocean circulation simulations. ODS and FL analyzed quantitative imaging data. TOD, PF and OJ analyzed metagenome-assembled genome data. GR, NH, AF-G, S Suweis, RN, J-MA, MM and EP contributed additional analysis. S Sunagawa, LG, PB, CB, MBS and EK provided additional interpretation of results. KL, EM and JP coordinated the genomic sequencing with the informatics assistance of CD, FG and J-MA. S Roux, JRB and MBS contributed viral data, PB and S Sunagawa contributed bacterial data. CB, S Romac, NH, CdV and DJR analyzed eukaryotic metabarcoding data. CD, SK, MP, S Searson and JP coordinated collection and management of *Tara* Oceans samples. *Tara* Oceans Coordinators provided support and guidance throughout the study. All authors discussed the results and commented on the manuscript.

## Author Information

The authors declare that all data reported herein are fully and freely available from the date of publication, with no restrictions, and that all of the samples, analyses, publications, and ownership of data are free from legal entanglement or restriction of any sort by the various nations in whose waters the *Tara* Oceans expedition sampled. Metagenomic and metabarcoding sequencing reads have been deposited at the European Nucleotide Archive under accession numbers provided in Supplementary Table 1. Contextual metadata of *Tara* Oceans stations are available in Supplementary Table 2. Metagenomic dissimilarity, OTU community dissimilarity, imaging community dissimilarity, simulated travel times, geographic distances and MAG dissimilarity are provided in Supplementary Tables 3-18. All Supplementary Tables, in addition to tables of 18S V9 barcodes and OTUs and the V9 reference database are available on FigShare at the following URL: https://doi.org/10.6084/m9.figshare.11303177 The authors declare no competing financial interests.

Correspondence and requests for materials should be addressed to Olivier Jaillon, Daniele Iudicone, Maurizio Ribero d’Alcalà, Colomban de Vargas.

## Data availability

https://doi.org/10.6084/m9.figshare.11303177 Supplemental Tables 1-21 (including DDBJ/ENA/GenBank short read archive identifiers for *Tara* Oceans metagenomic & 18S V9 sequence reads, and distance matrices), Datasets 1-3 (18S V9 metabarcode and OTU tables, and reference database).

## Supplementary Information

### Supplementary Information 1. Comparison of metagenomes and OTUs

Metagenomic comparisons reflect fine-scale differences in genome content at the community level as a function of diversity, genome size and organismal abundance, and also depend on the rate of evolution of each specific lineage. With exhaustive sampling, metagenomic dissimilarity could theoretically distinguish among genomes in a sample separated by a single mutation. However, our metagenomic sequencing level was likely not able to reach saturation due to the number of genomes per sample and their putative large size (metatranscriptomes, which contain fewer sequences per species than do metagenomes, did not reach saturation within *Tara* Oceans samples^59^). For example, if for a pair of samples we sequence 50% of the total amount of the unique genomic DNA present, we expect the maximum similarity of the two samples to be roughly 25% (0.5 x 0.5). Therefore, the pairwise metagenomic dissimilarities we calculated between samples probably reflected a combination of genomic differences weighted towards more abundant organisms. In contrast, OTUs, obtained by sequencing single marker genes, approach biodiversity saturation^8,18,19^. However, OTU resolution depends on the choice of the marker to be used, the threshold of similarity for the marker, and its lineage-specific substitution rate, and may therefore confound evolutionarily and/or ecologically distant organisms^60-64^. We observed a significant agreement between the two proxies (Supplementary Fig. 2), although dissimilarities based on OTUs were generally lower than those computed from metagenomic data (Supplementary Fig. 3a).

Analyses of plankton biogeography produced consistent results based on metagenomic and OTU data (Supplementary Fig. 4, Supplementary Fig. 5, Supplementary Fig. 8, Supplementary Fig. 9). For simplicity, in the main text, we chose to highlight results based on metagenomes rather than on OTUs for three reasons. First, the metagenomic sequencing protocol and subsequent measurement of dissimilarity was uniform across size fractions, whereas OTUs were defined differently for the viral-enriched, bacterial-enriched and eukaryote-enriched size fractions (Methods). Second, the biogeographical patterns we obtained (see below) may be more evident in comparisons among metagenomic sequences (our data source in identifying genomic provinces), as genomes accumulate single-base changes and other variants more quickly than a single ribosomal gene marker. Third, β-diversity estimated by metagenomic dissimilarity generally displayed higher correlation values with minimum travel time (T_min_; Supplementary Fig. 8).

### Supplementary Information 2. Robustness of genomic provinces

We assessed the robustness of genomic provinces in five separate ways. First, we tested 5 different hierarchical clustering algorithms from the R-package pvclust_1.3-2^45^ (UPGMA - Unweighted Pair Group Method with Arithmetic mean; McQuitty’s method; Complete linkage; Ward’s method; Single linkage) on the metagenomic pairwise dissimilarities produced by Simka separately for the six organismal size fractions, followed by multiscale bootstrap resampling. We used the cophenetic correlation coefficient from the R-package dendextend_1.5.2^65^ to measure how accurately the dendrograms produced by each method preserved the pairwise distances within the input dissimilarity matrices^66,67^. The ranking of the cophenetic correlation coefficient for different clustering methods within each size fraction (Supplementary Table 19) was consistent with a published large-scale methodological comparison of clustering methods for biogeography, which considered UPGMA agglomerative hierarchical clustering to have consistently the best performance^44^. Second, we compared clustering results among all size fractions using Baker’s Gamma Index^68^ from the R-package corrplot_0.77^69^, which is a measure of association (similarity) between two trees based on hierarchical clustering (dendrograms). The Baker’s Gamma Index is defined as the rank correlation between the stages at which pairs of objects combine in each of the two trees. For each type of correlation, the UPGMA was consistently the most correlated with other clustering methods (Supplementary Table 20). This allowed us to conclude, in agreement with previous results^44^, that the UPGMA method is likely more robust than the other methods we tested.

Third, we compared the genomic provinces found by our UPGMA hierarchical clustering approach to those found by two different non-hierarchical methods: K-means on the positions found by multidimensional scaling and spectral clustering on the nearest-neighbor graph. Both methods rely on (i) a dissimilarity matrix and (ii) a tuning parameter (dimension of the projection space for K-means, and number of neighbors for spectral clustering). K-means uses the numeric values of the dissimilarities, whereas spectral relies only on their ordering (e.g., community A is closer to B than to C). We compared the genomic provinces to clusters found by K-means and spectral clustering for all values of the tuning parameter using the Rand Index (RI; from the GARI function of the loe R package version 1.1^70^), a score of agreement between partitions. Results are reported as mean +/- s.d. of the RI: 1 means perfect agreement and 0 complete disagreement. Fourth, in order to assess the significance of the genomic provinces, we performed a multivariate ANOVA to partition metagenomic dissimilarity across regions, using the adonis function of the vegan R package version 2.5-4^42^. Note, however, that since the same data were used both to construct the genomic provinces and to assess their significance, the p-values estimated by ADONIS might be anti-conservative. The results of the third and fourth analyses are presented in Supplementary Table 21.

Fifth, we found that clustering of samples in genomic provinces was consistent with a complementary visualization based on the same data: RGB colors derived from the first three axes of a principal coordinates analysis (PCoA-RGB) of β-diversity, in which similar colors represent similar communities (Supplementary Fig. 4; see Methods). Samples within the same genomic province generally shared the same range of PCoA-RGB colors. Because the clustering approach was hierarchical, samples sharing some similarity could have been assigned to different genomic provinces due to binary decisions during the clustering process. This was also reflected in the PCoA-RGB colors, where the boundaries of genomic provinces did not indicate a complete change of communities among genomic provinces (and, conversely, belonging to the same genomic province did not imply identical communities). Nonetheless, samples with similar PCoA-RGB colors were generally situated in closely-related branches in the UPGMA tree (Supplementary Fig. 6). An illustrative example is genomic province F5 (of the 180-2000 μm size fraction; Supplementary Fig. 4f), which encompassed stations in the Atlantic, Mediterranean Sea and some subtropical stations in the Indo-Pacific. In this wide region, the PCoA-RGB colors indicate the variation in community composition within the genomic province, and also reflect the relatedness of F5 to its adjacent samples, in particular those in the subtropical Atlantic/Pacific region F4, its neighbor in the UPGMA tree (Supplementary Fig. 6f).

### Supplementary Information 3. Comparison of genomic provinces to previous biogeographical divisions

Current approaches in biogeographic theory divide the ocean into regions based either on expert knowledge applied to satellite data, as in the hierarchical nesting by Longhurst^11^ into biomes (macro-scale, essentially representing a division of the world’s oceans into cold and warm waters, and coastal upwelling zones) and biogeochemical provinces (BGCPs, areas within biomes defined by observable boundaries and predicted ecological characteristics), or, alternatively, into the objective provinces of Oliver and Irwin^53^, which are based solely on statistical analyses. Longhurst BGCPs are based upon, primarily, monthly variations of chlorophyll *a*, the geography of the seasonal cycle of physical factors (such as the depth of the upper ocean mixed layer) and surface temperatures. In turn, these ocean properties are strongly modulated by oceanic currents (for example, moderate to large mixed layer depths are observed generally on the poleward side of the subtropical gyres). In contrast, the objective global ocean biogeographic provinces proposed by Oliver and Irwin^53^ were based upon clustering temporal variability of chlorophyll concentration and surface temperatures, both measured from satellite data. They combined a proxy for the intensity of primary productivity with water temperature, therefore emphasizing regions similar in their temporal variability for both properties (which essentially corresponds to the seasonal cycle). None of these ocean partitionings directly considered organismal community composition.

We tested whether genomic provinces were comparable with these partitionings by performing an analysis of similarity (ANOSIM; Supplementary Fig. 4a-f, insets; Methods). The four small size classes, 0-0.2 μm, 0.22-1.6/3 μm, 0.8-5 μm, and 5-20 μm (Supplementary Fig. 4a-d) were more consistent with Longhurst BGCPs. In contrast, for the two larger size fractions 20-180 μm and 180-2000 μm, the three biogeographical divisions were not strongly different within the ANOSIM (Supplementary Fig. 4e-f).

From an oceanographic point of view, plankton should be quasi-neutrally redistributed (i.e., homogenized) by currents and their biogeography should follow the structure of the main recirculations, within a range of physiologically compatible temperatures. In this point of view, our results are consistent with the large-scale geographic distributions found by Hellweger *et al.*^14^ using a neutral model.

### Supplementary Information 4. Differences in genomic province sizes among organismal size fractions

Globally, we obtained more numerous, smaller genomic provinces in the smaller size fractions and fewer, larger genomic provinces in the larger size fractions (Supplementary Fig. 4, Supplementary Fig. 7). We observed a similar pattern using OTU data (Supplementary Fig. 5). Whereas smaller size fractions generally lacked geographically widespread genomic provinces containing numerous *Tara* Oceans samples, the two largest size fractions were both characterized by two very widespread genomic provinces: F5 and F8 for the 180-2000 μm size fraction, and E5 and E6 for the 20-180 μm size fraction. These large genomic provinces were latitudinally limited by the boundary between the subtropics and subpolar regions, and spanned different oceanic basins. Notably, in the Southern Hemisphere the subtropical gyres actually form a single supergyre^71^ and there are almost no metabolic (mainly temperature) barriers between the northern and southern subtropical gyres (see Supplementary Fig. 4), potentially explaining genomic provinces in the 20-180 μm and 180-2000 μm size fraction that contain samples from the North and South Atlantic. For example, in the 180-2000 μm size fraction, F5 mostly covered the North and South Atlantic Oceans and adjacent systems, and F8 covered the Indo-Pacific low- and mid-latitudes. No clear correspondence existed with biogeochemical patterns (e.g., nutrient ratios), except for the clusters coinciding with upwelling systems (F3 for the California upwelling, F7 for the Chile-Peru upwelling and F2 for the Benguela upwelling system) and for the samples collected at the deep chlorophyll maximum (DCM) in the Pacific subtropical gyres (F5); this is consistent with the comparison of genomic provinces to previous biographical divisions, in which the genomic provinces of smaller size fractions were more consistent with Longhurst BGCPs, but those of larger size fractions were not (Supplementary Information 3). A bimodal zooplankton species distribution (split into subtropical and subpolar communities, with ubiquitous warm water species) was also detected by a recent study on copepod population dynamics that used alternative approaches to analyze the same metagenomic dataset^72^ (see their Fig. 2). More locally, within the North Atlantic (see also Supplementary Information 6), along the northern boundary of the subtropical gyre, cold and warm copepod species overlapped because of cross-current dispersal. Nonetheless, although both cold and warm species appeared to be able to travel long distances, mixing among them was not sufficient to create a local genomic province in our data.

We interpret the difference in genomic province sizes between smaller and larger size fractions as the result of various factors. Plankton smaller than 20 μm (femto-, pico- and nanoplankton), which represent most of the prokaryotic and eukaryotic phototrophs^18,19^, are sensitive to a suite of environmental factors (i.e., temperature^73^, nutrients and trace elements^2^; see also Supplementary Fig. 7) and generally have a shorter life cycle, together leading to faster fluctuations in their relative abundance in the communities we sampled. In contrast, larger plankton have longer life cycles and, if they are predators that are not strongly selective in their feeding, or are photosymbiotic hosts capable of partnering with multiple different symbionts, may cope with local fluctuations in environmental conditions. Therefore, they should be affected primarily by large scale, mostly latitudinal, variations in the environment, leading to larger genomic provinces, whereas smaller plankton are grouped into smaller provinces more influenced by local environmental conditions. Overall, this difference in biogeography suggests a size-based decoupling between smaller and larger plankton (which may also extend to nekton such as tuna and billfish^74^), with implications for the structure and function of oceanic food webs and other types of biotic interactions.

### Supplementary Information 5. Genomic provinces as stable ecological continua

As plankton communities are transported by ocean currents, they change over time due to the various processes that occur in the context of the seascape: variations in temperature, light and nutrients (where changes in the latter may also be induced by plankton communities), intra- and inter-individual and species biological interactions, and mixing with neighboring water masses. Thus, a continuum of composition among nearby samples is expected as a natural consequence of community turnover within the seascape over time. We observed the effects of continuous turnover in our biogeographical analyses (Fig. 1a, Supplementary Fig. 4, Supplementary Fig. 5, Supplementary Information 2) in which nearby samples often reflected gradual, but not complete changes in community composition.

We measured the time window of transport by currents separating two samples during which the changes in their community composition were maximally correlated with travel time, resulting in a global average of T_min_ < roughly 1.5 years. This represents the travel time during which predictable continuous turnover occurs in our dataset. Notably, T_min_ does not necessarily define the turnover rate itself, which depends on how strongly different seascape processes affect communities with differing biological characteristics (see Supplementary Information 6).

The global ocean current system is composed of a series of large-scale main currents and associated recirculations (which are also referred to as gyres). Therefore, we present the following hypothesis as a potential explanation of our results: the average global timescale of 1.5 years is comparable to the crossing time of an ocean gyre (i.e., the amount of time it takes a water parcel to travel from one side of a gyre to the other), e.g., to cross the North Atlantic basin while riding the Gulf Stream system. This time scale of 1.5 years is probably an underestimate, since our sparse sampling did not cover all current systems. Within different systems, the transport by main currents leads to stable, continuous patterns of changes in community structure and nutrient concentrations, and also explains how temporally stable genomic provinces can exist in the face of ocean circulation. Within each system we have thus to expect that community turnover is long enough to allow for this long range predictability due to smooth, continuous changes. Significant heterogeneity in environmental conditions among different circulation patterns means that moving from system to another (and therefore, in our case here, beyond the 1.5 year timescale; Supplementary Fig. 9c-f) disrupts the interlinked relationship among local seascape processes, leading to a global delimitation into separate ecological continua among different gyre-scale current systems.

### Supplementary Information 6. Community turnover in the North Atlantic

In order to characterize the impact of physical and biological processes on changes in metagenomic composition during travel along currents, we focused on the well-known current systems crossing the North Atlantic into the Mediterranean Sea (the Gulf Stream and other currents around the subtropical gyre^22,75-77^; Supplementary Fig. 10a). Across this region, the piconanoplankton (0.8-5 μm) were split into three genomic provinces, C5, C8 and C3, each less than 5,000 km wide (~11 months of travel time; Supplementary Fig. 4c). In contrast, mesoplankton (180-2000 μm) biogeography corresponded to a single province, F5, spanning from the Caribbean to Cyprus (> 9,700 km or ~18 months of travel time; Supplementary Fig. 4f; see also Supplementary Information 4). Metagenomic dissimilarity and T_min_ were strongly correlated within the region (Spearman’s ρ between 0.44 and 0.86 depending on size fraction, Supplementary Fig. 10b-e), which allowed us to explore the relationship of genomic province size, ocean transport and plankton community turnover over scales from months to years. We calculated metagenomic turnover times as e-folding times based on an exponential fit of metagenomic dissimilarity to T_min_ (ranging from a few months to a few years, Methods). The metagenomic turnover time of smaller plankton (< 20 μm) was approximately one year. In contrast, for the larger size fractions, the metagenomic turnover time was approximately two years, suggesting that a lower turnover rate for larger plankton may explain their geographically larger genomic provinces.

We note that our results on metagenomic turnover time appear different from a recently published study that also calculated turnover rates for plankton, which found faster rates for larger organisms^15^. This may be explained by two significant differences between our approach and theirs: first, their measurements of β-diversity were based on presence/absence (Jaccard) comparisons among either morphological species or OTUs, whereas our calculations of turnover time above were based on metagenomic sequences. As described above (Supplementary Information 1), there are significant differences in resolution between OTU-based and metagenomic data, and we would expect similar differences in resolution between organismal observation data and metagenomic sequences. In fact, due to these differences in resolution, our estimates of metagenomic turnover based on OTU rather than metagenomic data show a similar trend to those of Villarino *et al.*^15^ (Supplementary Fig. 10f-i). Second, their turnover rates were calculated separately for individual plankton groups (the 9 main groups were prokaryotes, coccolithophores, dinoflagellates, diatoms, all microbial eukaryotes, gelatinous zooplankton, mesozooplankton, macrozooplankton and myctophids), whereas our metagenomic data represent samples of the full plankton community within each size fraction. Among these, several groups (e.g., dinoflagellates or mesozooplankton) would be expected to be found across multiple *Tara* Oceans size fractions, blurring potential comparisons. Thus, our study and Villarino *et al.* calculated rates of change using broadly similar approaches, but based on very different underlying biological substrates.

### Supplementary Information 7. Plankton biogeography is robust to missing samples

Although many individual *Tara* Oceans stations are missing metagenomic or metabarcode sequencing data for a subset of size fractions (Supplementary Fig. 1b), all oceanic regions have broad coverage for each size fraction, with the exception of the viral-enriched size fraction in the North Atlantic. In fact, the largest source of missing data in our study is due to limited sampling of the viral-enriched size fraction in this region. Nevertheless, we found a pattern for organismal biogeography and for its relation with transport time that is not dependent on the size fraction, and therefore also does not depend on the particular size fractions sampled at specific set of sampling sites. In our analyses, we found a consistently similar patterns across the 4 smaller size fractions (each fraction was sampled and analyzed independently from the others) as opposed to the two larger ones. In addition, for our results relating to ocean transport time, the fact that the sampling sites are not exactly the same among size fractions actually lends robustness to our results, since it means that the dynamics we found are not overly dependent on any one selected site, region, or subset of sampling stations.

---

**Supplementary Figure 1.**
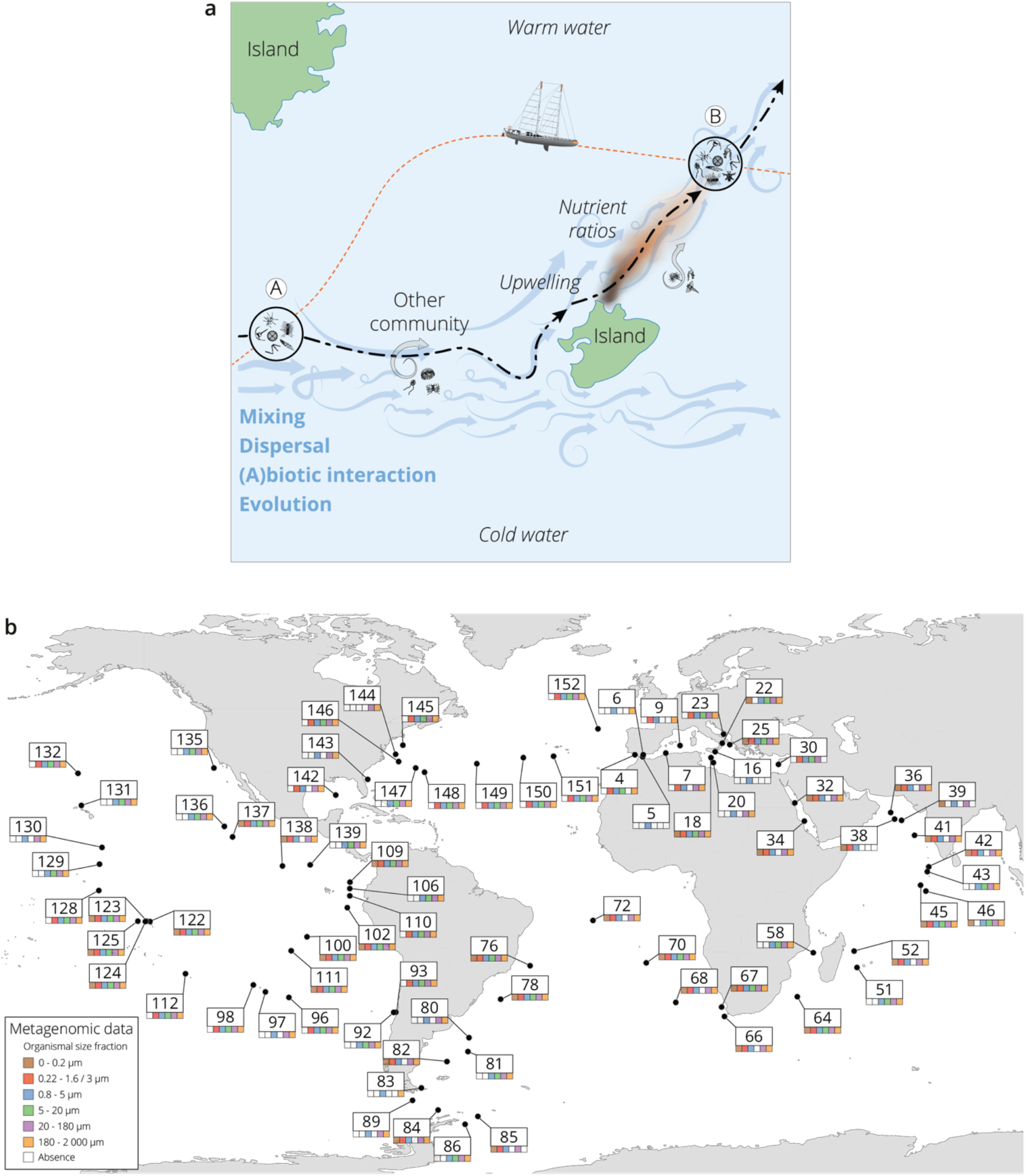
The seascape, plankton transport and community metagenomic samples of *Tara* Oceans stations. **a**, A community sampled at a given location (A) changes over time as it travels along ocean currents (dashed bold line) to a second location (B). It is affected by numerous external processes, including mixing with water containing other communities and changes in local nutrient concentration, and by internal processes, such as biotic interactions. In this study, the *Tara* schooner followed a sampling route (orange dashed line) leading to an elapsed time between the 2 sampling sites A and B that was independent of plankton travel time. **b**, Location, station number, and sequenced surface metagenomic samples.

**Supplementary Figure 2.**
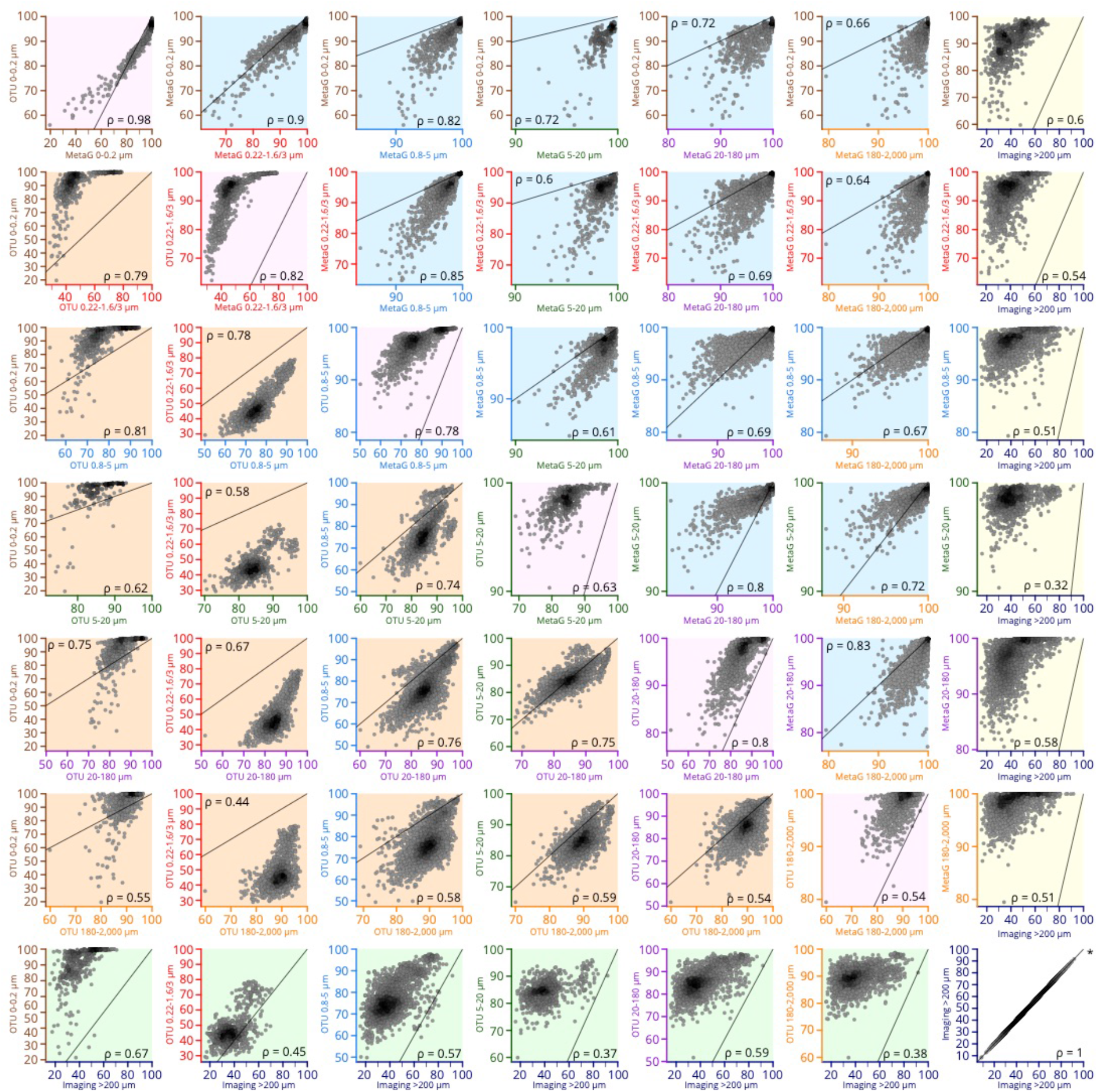
Scatter plots comparing β-diversity estimates from metagenomic, OTU-based and imaging-based dissimilarity. Source data for comparisons are indicated on the axes of each plot (axis colors correspond to size fractions or imaging data as in other figures, e.g., Supplementary Fig. 9). Axes are not necessarily drawn on the same scales; the identity line is indicated on each plot to help interpret the relationship between axes. Plots with a pink background are comparisons of metagenomic versus OTU-based dissimilarity within the same size fraction. Plots with a blue background are comparisons of metagenomic dissimilarity among size fractions, and those with an orange background compare OTU-based dissimilarity among size fractions. Plots with a yellow or green background compare imaging-based dissimilarity to either metagenomic or OTU-based dissimilarity, respectively. Each point within a plot represents a pairwise comparison of β-diversity estimates between two *Tara* Oceans samples. Rank-based correlations (Spearman, p ≤ 10^-4^) are indicated in each plot.

**Supplementary Figure 3.**
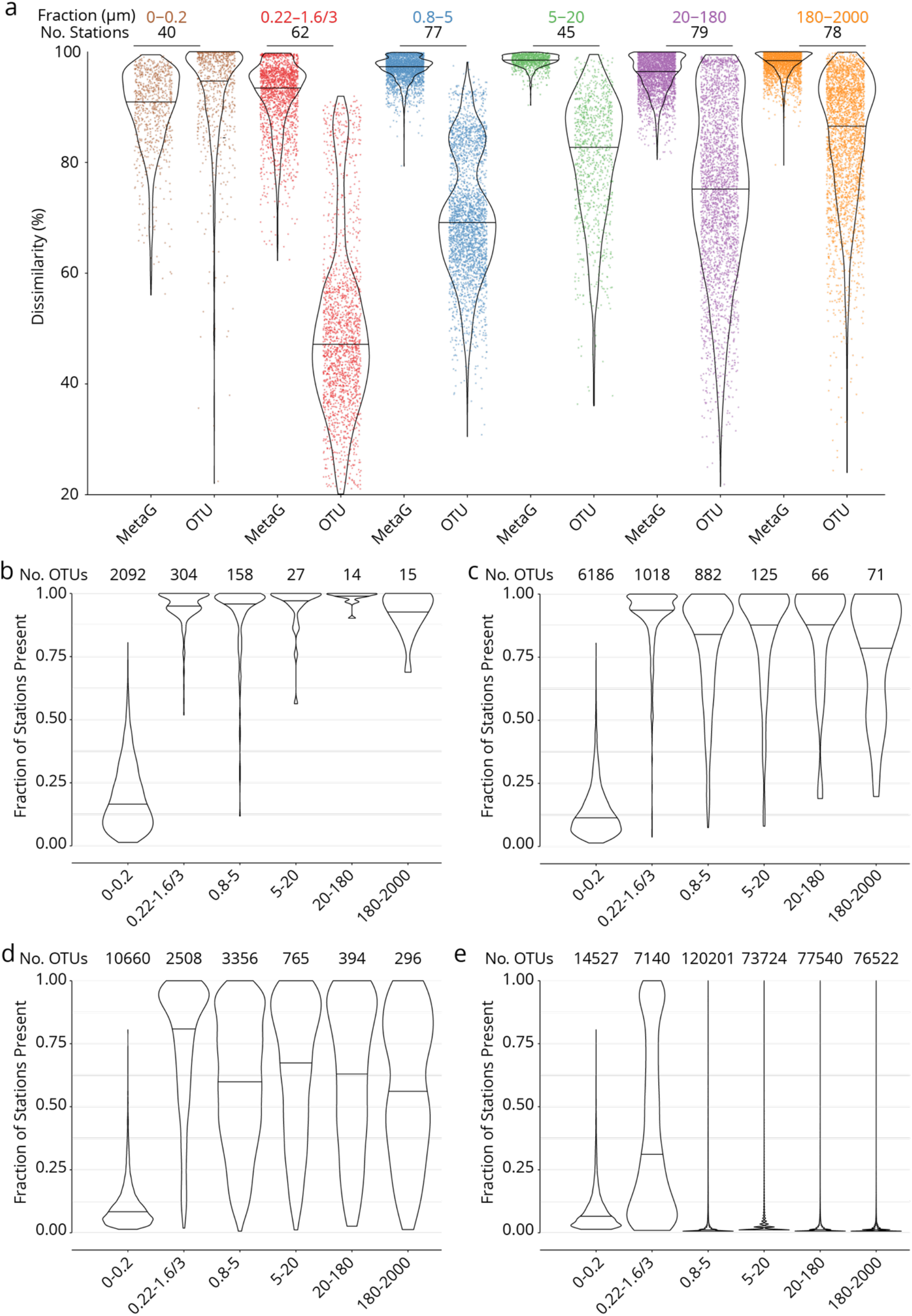
Global dissimilarity and OTU occupancy. **a**, Distributions of dissimilarity for six organismal size fractions (measured either as metagenomic or OTU dissimilarity; see Supplementary Information 1). One colored point represents one pair of stations. Violin plots (horizontal line: median) summarize each distribution. The number of stations in common between the metagenomic/OTU data sets within each size fraction is indicated above. **b-e**, **OTU occupancy for different proportions of total abundance.** Fraction of stations present (occupancy) for the minimum number of OTUs (indicated above) necessary to represent different proportions of the total abundance within each organismal size fraction. A relatively small number of abundant and cosmopolitan taxa represents the majority of the abundance within each size fraction; this effect is more pronounced with increasing organismal size. **b**, OTUs representing 50% of the total abundance within each size fraction. **c**, 80%. **d**, 95%. **e**, 100% (all OTUs).

**Supplementary Figure 4.**
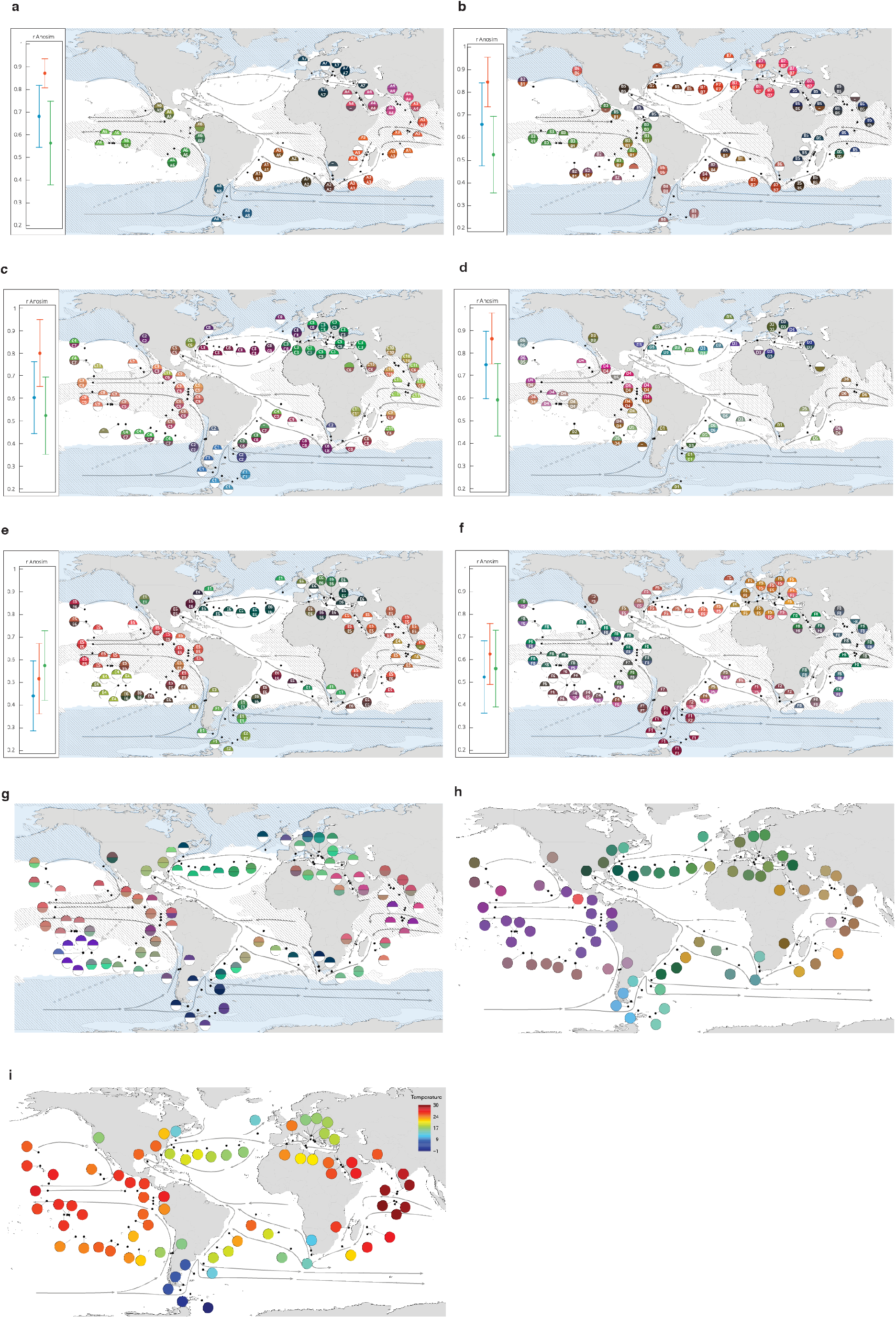
Genomic provinces in comparison to previous ocean divisions and to metagenome assembled genome abundance variation, and ordination maps of environmental parameters. Colors are based on PCoA-RGB (Methods) and do not correspond directly among maps. **a-f**, Geographical maps of genomic provinces by organismal size fraction (see Supplementary Information 2). Circles denote stations with data available for the size fraction and contain the corresponding genomic province identifiers (one letter prefix per size fraction (A-F); stations not assigned to genomic provinces are shown as ‘-’). The top portion of each circle represents samples collected at the surface and the bottom portion represents the deep chlorophyll maximum (stations missing metagenomic data for one of the two depths are drawn as half circles). Major currents are shown with solid black arrows, wind transport with dashed grey arrows. Blue zones indicate temperature < 14 °C. Hashed zones indicate phosphate concentration > 0.4 mmol. Hierarchical dendrograms that were used to build genomic provinces are shown in Supplementary Fig. 6. Maps with colors based on OTU dissimilarity are shown in Supplementary Fig. 5. **a**, ‘A’ prefix, 0-0.2 μm size fraction. **b**, ‘B’ prefix, 0.22-1.6/3 μm. **c**, ‘C’ prefix, 0.8-5 μm. **d**, ‘D’ prefix, 5-20 μm. **e**, ‘E’ prefix, 20-180 μm. **f**, ‘F’ prefix, 180-2000. **Insets**, Results of ANOSIM to determine, independently for each size fraction, the ability of three nested levels of ocean partitioning to explain metagenomic dissimilarities among stations (blue, Longhurst biomes; red, Longhurst biogeochemical provinces; green, Oliver and Irwin objective provinces; see Methods and Supplementary Information 3). **g**, Geographical map for the 20-180 μm size fraction, for comparison with panel **e**, generated from metagenome assembled genome (MAG) dissimilarity among stations. **h**, The distribution of temperature and nutrient variations matches the biogeography of small plankton (< 20 μm). Stations are colored based on an ordination of Euclidean distances in temperature, NO_3_ + NO_2_, PO_4_ and Fe. **i**, The distribution of temperature matches the biogeography of large plankton (> 20 μm). Stations are colored following a Box-Cox transformation (Methods).

**Supplementary Figure 5.**
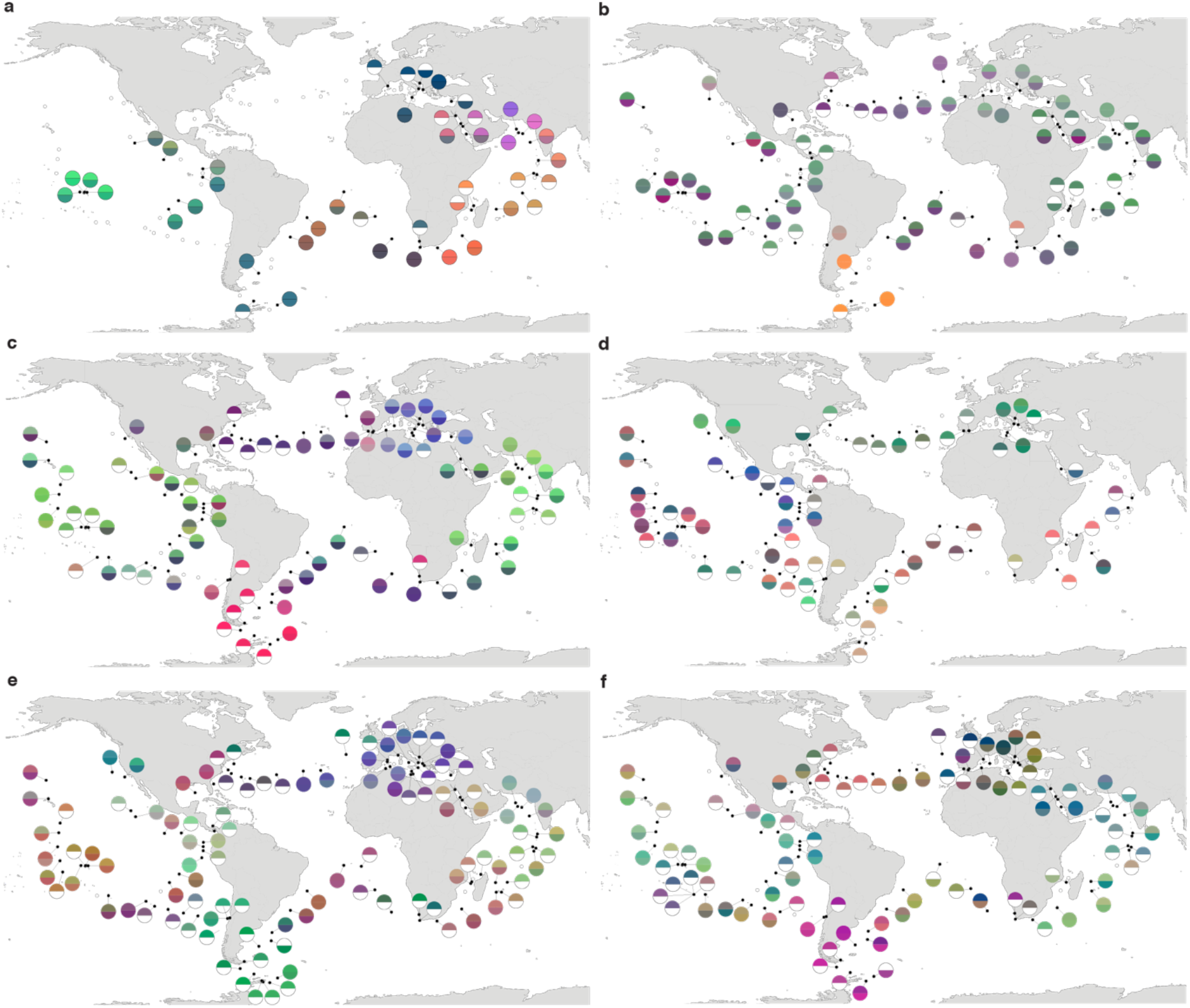
Biogeography based on an ordination of OTU dissimilarity. **a-f**, Principal coordinates analysis (PCoA)-RGB color maps for OTUs (see Methods). The top of each half circle represents samples collected at the surface and the bottom portion represents the deep chlorophyll maximum (stations missing OTU data for one of the two depths are drawn as half circles). Station colors do not correspond among size fractions. **a**, 0-0.2 μm size fraction. **b**, 0.22-1.6/3 μm. **c**, 0.8-5 μm. **d**, 5-20 μm. **e**, 20-180 μm. **f**, 180-2000 μm.

**Supplementary Figure 6.**
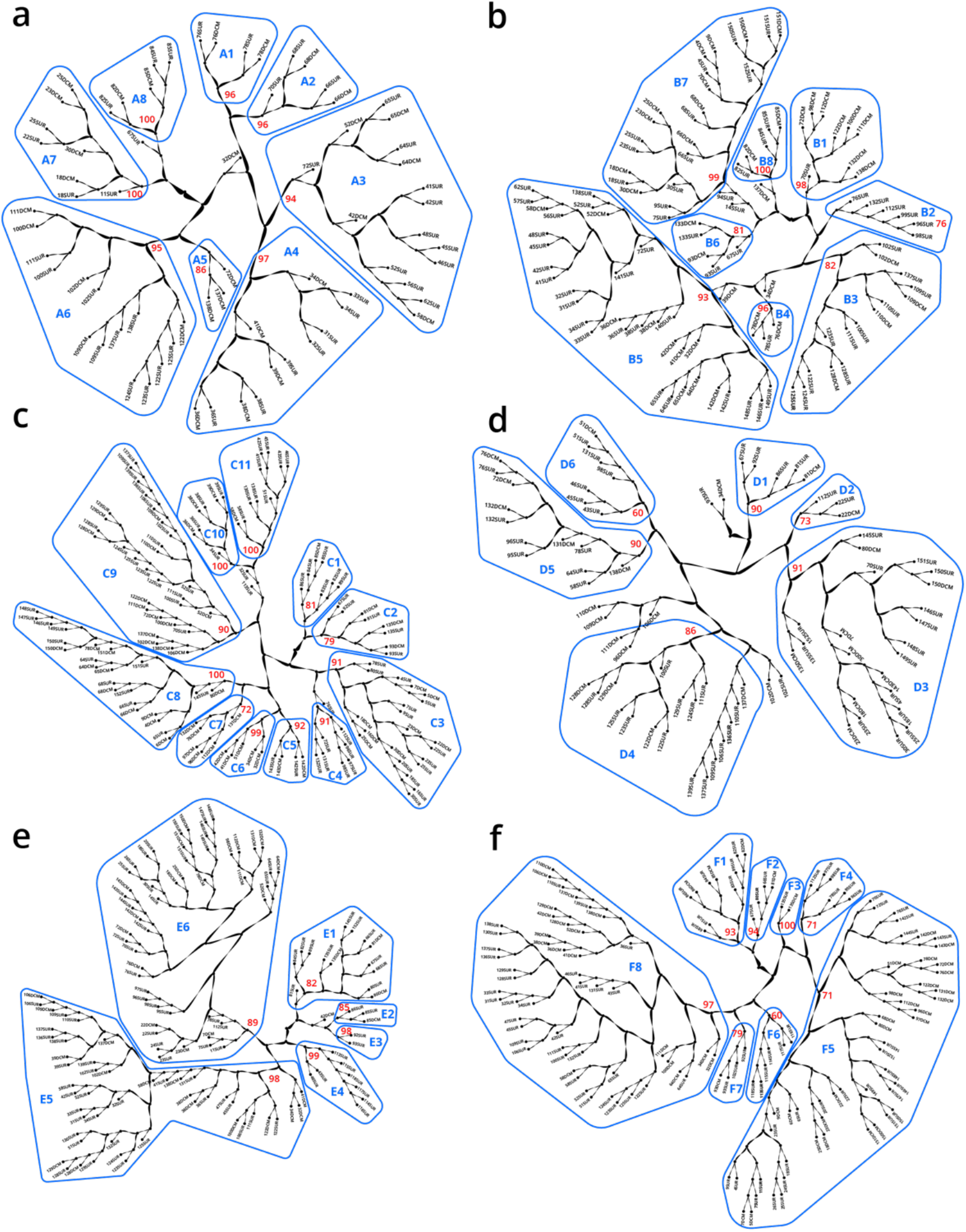
Hierarchical trees illustrating how samples were partitioned into genomic provinces. Dendrograms resulted from UPGMA clustering. Each sample (SUR: surface, DCM: deep chlorophyll maximum) is shown as a leaf. Genomic provinces are shown with their identifiers in blue polygons; identifiers are composed of one letter prefix per size fraction (A-F) and a number. Bootstrap values in red show the support at the key nodes that separate genomic provinces from one another. See also Supplementary Information 2 on the robustness of genomic provinces. **a**, ‘A’ prefix, 0-0.2 μm size fraction. **b**, ‘B’ prefix, 0.22-1.6/3 μm. **c**, ‘C’ prefix, 0.8-5 μm. **d**, ‘D’ prefix, 5-20 μm. **e**, ‘E’ prefix, 20-180 μm. **f**, ‘F’ prefix, 180-2000 μm.

**Supplementary Figure 7.**
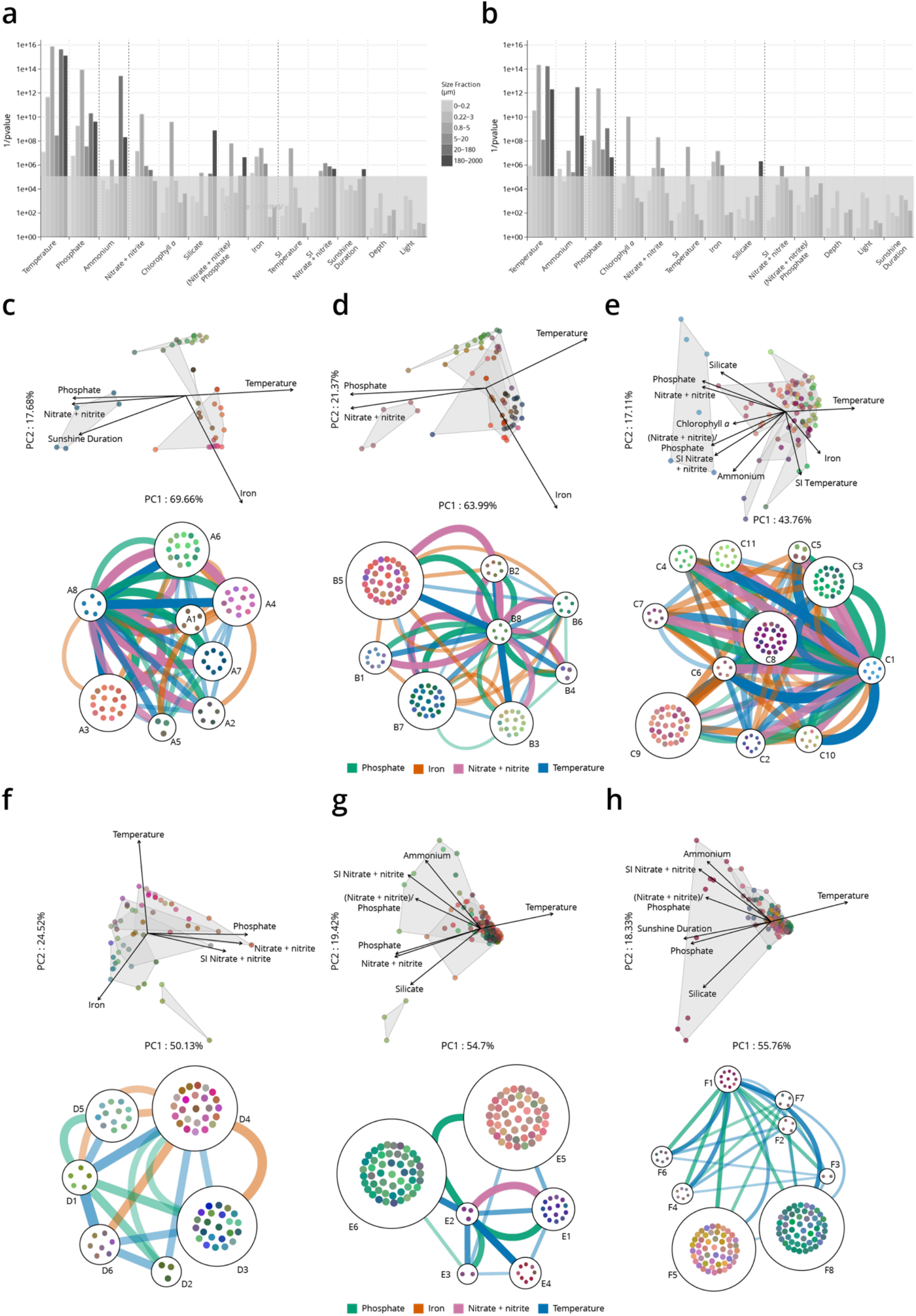
Environmental parameters that distinguish genomic provinces. **a-b**, Environmental parameters that significantly differentiate among genomic provinces (Kruskal-Wallis test, grey box indicates p values > 10^-5^). SI = Seasonality Index. **a**, all stations. **b**, Antarctic stations removed (see Methods). Eliminating Antarctic stations does not result in a large change in the parameters that significantly differentiate among provinces. **c-h**, Two types of visualizations of the relationships between genomic provinces and environmental parameters. Sample colors are those from Supplementary Fig. 4a-f. **Top plots within panels c-h**: principal components analysis-based visualization. Samples, and environmental parameters differing significantly (p ≤ 10^-5^) among genomic provinces, are projected onto the first two axes of variation. Grey polygons enclose different genomic provinces. **Bottom plots within panels c-h**: network-based visualization. Each genomic province is represented as a node, with the individual samples composing the province within the node. Edges between nodes represent differences in temperature, nitrate + nitrite, phosphate and iron that significantly differentiate (p ≤ 10^-5^) among genomic provinces, that are statistically significantly different between individual pairs of genomic provinces (*post hoc* Tukey test, p < 0.01) and whose difference in median parameter values is ≥ 1 standard deviation (calculated from the parameter values of all samples in the size fraction). Thicker edges represent larger differences. **c**, 0-0.2 μm size fraction. **d**, 0.22-1.6/3 μm. **e**, 0.8-5 μm. **f**, 5-20 μm. **g**, 20-180 μm. **h**, 180-2000 μm.

**Supplementary Figure 8.**
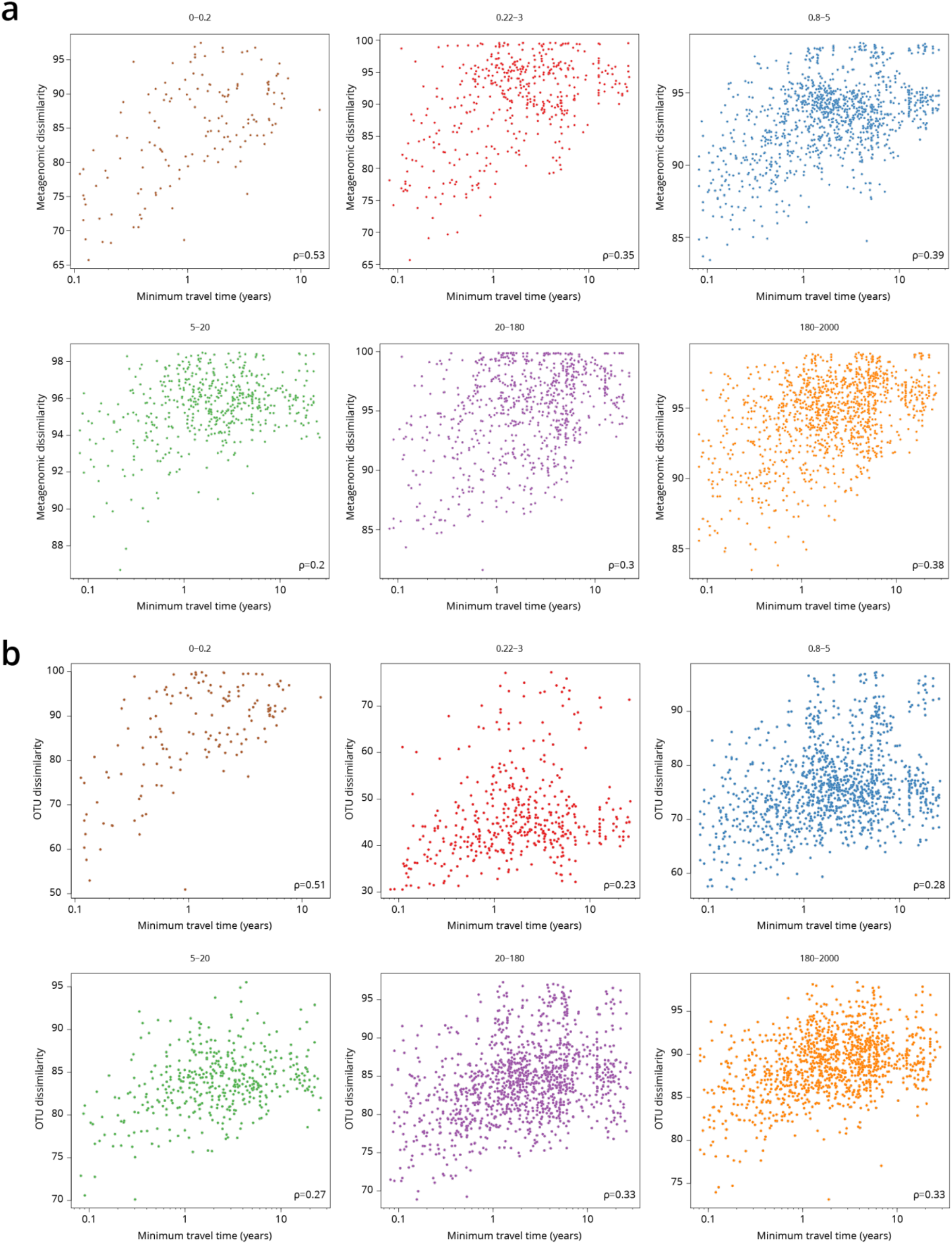
Global correlations of dissimilarity with minimum travel time (T_min_). Scatter plots of dissimilarity versus T_min_. One point represents a pair of samples. **a**, metagenomic dissimilarity. **b**, OTU dissimilarity. Global Spearman correlation values are indicated within each panel.

**Supplementary Figure 9.**
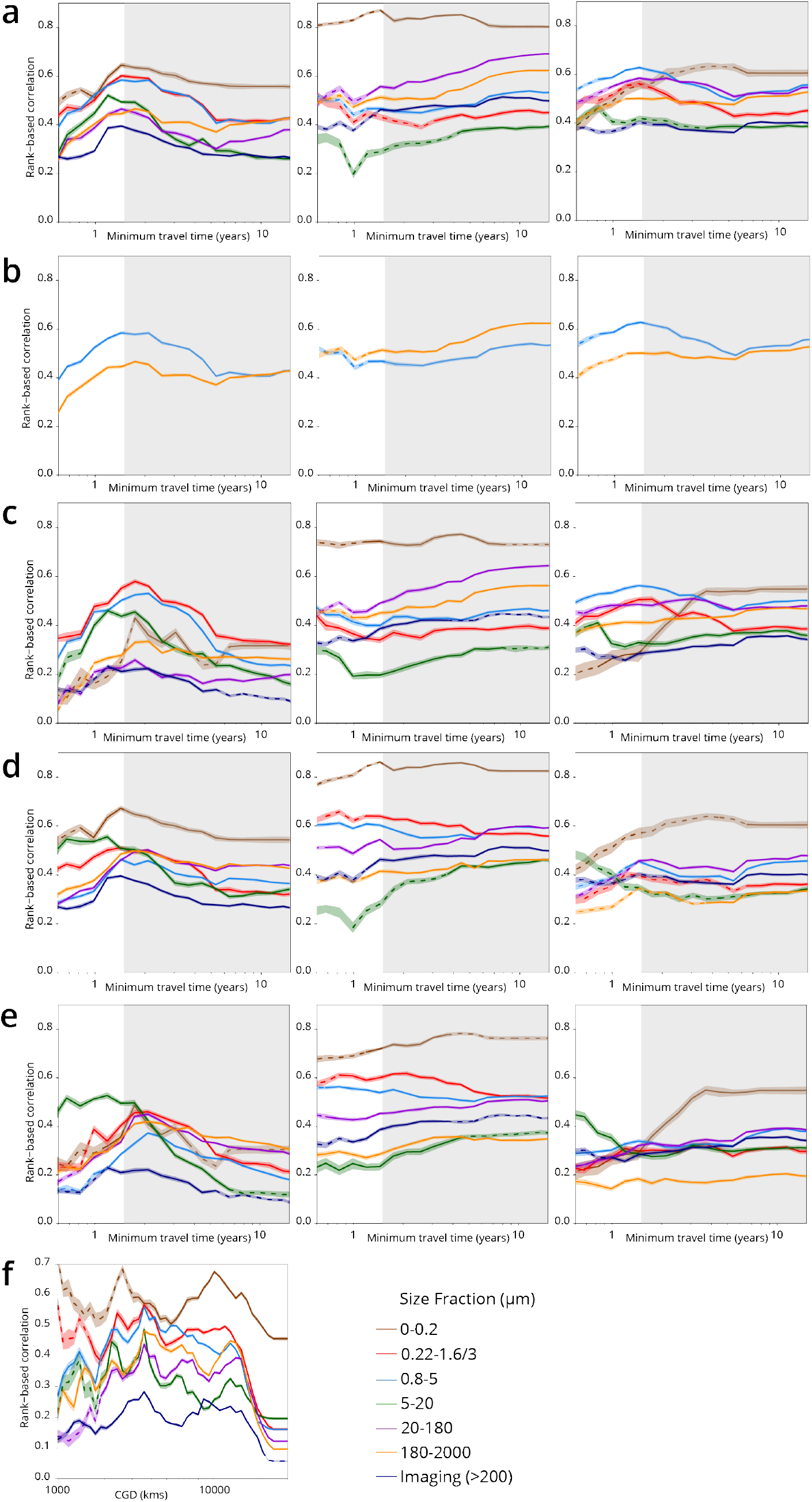
Plankton travel time, dissimilarity, environmental distance and geographic distance show different temporal patterns of pairwise correlation. Spearman correlation values are shown separately by organismal size fraction. Non-significant correlations (p > 0.01) are shown with dashed lines. **a-e**, Correlations for pairs of *Tara* Oceans samples separated by a minimum travel time less than the value of T_min_ on the x axis. T_min_ >1.5 years is shaded in grey. Left panels: correlation of dissimilarity with T_min_; middle panels, dissimilarity with temperature; right panels: dissimilarity with differences in NO_3_ + NO_2_, PO_4_ and Fe. **a-c**, metagenomic dissimilarity. **d-e**, OTU dissimilarity. Correlations for imaging dissimilarity are superimposed on plots in **a** and **c-e**, for comparison. There is a maximum correlation of dissimilarity with T_min_ (and, for most size fractions, of dissimilarity with nutrients) for T_min_ <~1.5 years, but the correlation between dissimilarity and temperature does not display a similar maximum. **b** displays only the 0.8-5 μm (blue) and 180-2000 μm (orange) size fractions from **a**, to highlight that for smaller plankton, correlations with differences in nutrient concentrations were stronger for T_min_ up to ~1.5 years, but for larger plankton, correlations were stronger with temperature variations for T_min_ beyond ~1.5 years. **c** and **e**, Partial correlations to estimate the independent effects of T_min_ and environmental distances on β-diversity. Left panels: controlling for differences in temperature and for differences in NO_3_ + NO_2_, PO_4_ and Fe; middle and right panels: controlling for T_min_. Partial correlations do not affect the maximum correlation of dissimilarity with T_min_ for T_min_ <~1.5 years. **f**, Correlation of geographic distance (without traversing land; CGD) with metagenomic dissimilarity or imaging dissimilarity for pairs of *Tara* Oceans samples separated by a geographic distance less than the value on the x axis.

**Supplementary Figure 10.**
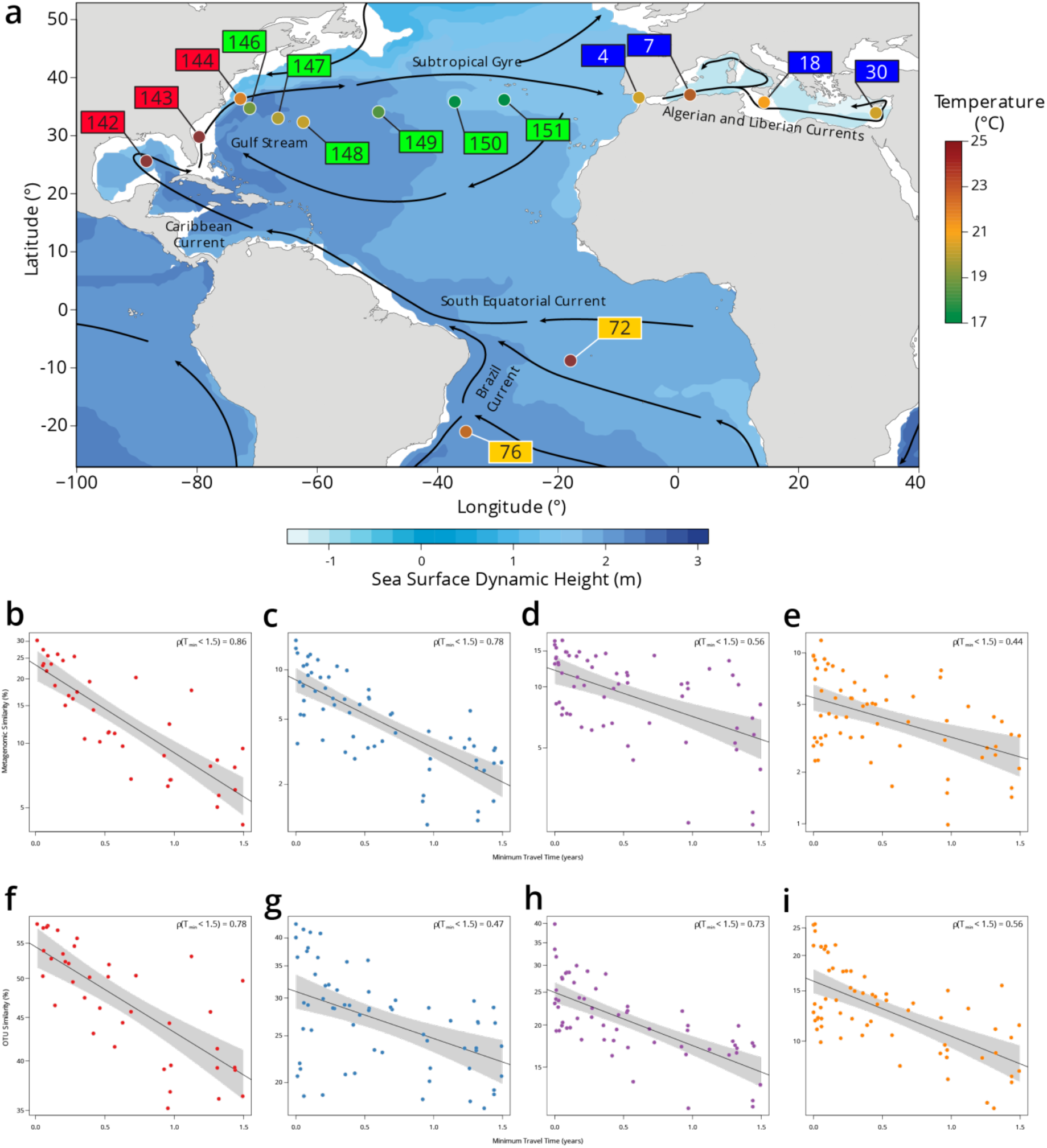
Plankton community composition turnover through the North Atlantic. **a**, Map of *Tara* Oceans stations, currents (solid lines), temperature by station (colored circles) and sea surface climatological dynamic height from CARS2009 (http://www.cmar.csiro.au/cars). Each station label has a color corresponding to a sub-region: South Atlantic in orange, Gulf Stream in red, Recirculation/Gyre in green and Mediterranean Sea in blue. **b-e**, Scatter plots of metagenomic similarity versus minimum travel time (T_min_) for these stations in the **b**, 0.22-3 μm; **c**, 0.8-5 μm; **d**, 20-180 μm; and **e**, 180-2000 μm size fractions. **f-i**, Scatter plots of OTU community similarity for the **f**, 0.22-3 μm; **g**, 0.8-5 μm; **h**, 20-180 μm; and **i**, 180-2000 μm size fractions. The black line represents an exponential fit, with a light grey shaded 95% confidence interval. The resulting turnover times using metagenomic similarity are τ = 0.91 y for 0.22-3 μm, τ = 0.91 y for 0.8-5 μm, τ = 2.22 y for 20-180 μm and τ = 1.99 y for 180-2000 μm. Turnover times using the OTU community similarity are τ = 4.23 y for 0.22-3 μm, τ = 4.08 y for 0.8-5 μm, τ = 2.6 y for 20-180 μm and τ = 2.1 y for 180-2000 μm. The viral-enriched 0-0.2 μm and the nanoplanktonic 5-20 μm size fractions are not shown due to insufficient sampling of these stations.

